# snRNA-seq analysis of the moss *Physcomitrium patens* reveals a conserved cytokinin-ESR module promoting pluripotent stem cell identity

**DOI:** 10.1101/2024.07.31.606100

**Authors:** Yuki Hata, Nicola Hetherington, Kai Battenberg, Atsuko Hirota, Aki Minoda, Makoto Hayashi, Junko Kyozuka

**Affiliations:** Grad. Sch. Life Sci. Tohoku Univ., Katahira 2-1-1 Aoba-ku, Sendai, Japan; RIMLS, Radboud University, Geert Grooteplein Zuid 10, 6525 GA, Nijmegen, Netherlands; University of Idaho, Stillinger Herbarium 875 Perimeter Dr. MS 3026, Moscow, ID 83844, USA; RIKEN CSRS, 1-7-22 Suehiro-cho, Tsurumi-ku, Yokohama, Japan

## Abstract

The shoot apical meristem (SAM), which contains pluripotent stem cells, serves as the source of the entire shoot system in land plants. To uncover the mechanisms underlying SAM development and its orign, we employed single-nucleus RNA sequencing technology in *Physcomitrium patens*, which contains a single stem cell known as the gametophore apical cell. We identified distinct cell clusters representing major cell types of the *P. patens* gametophyte, including the gametophore apical cells. We revealed dynamic gene expression changes during cell fate progression in the gametophore apical cell and found upregulation of cytokinin biosynthesis genes in this cell. We also identified *ENHANCER OF SHOOT REGENERATION 1* (*ESR1*) orthologs as key regulators of gametophore apical cells downstream of cytokinin. Given that ESRs promote SAM formation under cytokinin in angiosperms, we propose that the cytokinin-ESR module represents a conserved mechanism promoting stem cell identity that evolved in the common ancestor of land plants.

## Introduction

The terrestrialization of plants 500 million years ago marked a pivotal moment in the evolution of living organisms and the global environment.^1,2^ During this evolutionary event, land plants developed complex, three-dimensional structures suited for terrestrial habitats.^3–5^ A key advancement was the emergence of the apical meristem, which contains pluripotent stem cells that continuously supply cell populations capable of differentiating into various cell types.^2,6–9^ The shoot apical meristem (SAM) is the source of the entire the shoot system.^9–11^ The presence of SAM in two distinct developmental phases, i.e., the gametophyte (haploid) and sporophyte (diploid), suggests that the SAM evolved at least twice, each giveing rise to the meristem of gametophyte and sporophyte.^8,10^ Bryophytes and vascular plants are sister groups.^12^ While bryophytes possess the SAM in the gametophyte, vascular plants form the SAM in the sporophyte.^8,10^ Since green algae, the sister group of land plants, lack multicellular sporophytes, it is likely that SAM first evolved in the gametophyte and later in the sporophyte.^6,13^ Therefore, investigating bryophyte SAM provides an opportunity to uncover fundamental molecular mechanisms for stem cell specification common to both gametophyte and sporophyte SAMs.^10,14^ Additionally, it offers insights into the origin and diversification of plant stem cell systems. Intriguingly, the SAM of bryophytes consists of a single stem cell known as the ‘apical cell’, suggesting that cell fate determination is regulated at a single cell level.^10,14,15^ Therefore, elucidating the minimal mechanisms governing the pluripotency in the apical cell is of great interest.^10,16^

Reverse genetics studies using model bryophyte species have revealed the molecular basis for the diversification and conservation of the stem cell system.^10,14^ One of the important findings is that *KNOTTED-like HOMEOBOX I* (*KNOXI*) and *WUSCHEL-related HOMEOBOX* (*WOX*) family genes, which are crucial for stem cell maintenance in angiosperms, are not essential for bryophyte SAMs.^17,18^ On the other hand, auxin, cytokinin, and CLAVATA3/EMBRYO SURROUNDING REGION-RELATED (CLE) peptide signalings are critical for both bryophytes and angiosperm SAMs.^5,19–21^ Transcriptomic approaches also highlighted the similarities and differences between bryophytes and angiosperms SAMs.^13,22^ Recent progress in single-cell RNA-seq technologies has facilitated the characterization of gene expression profiles at single-cell resolution.^23^ This technology has been employed successfully across various plant tissues, including the SAM, root apical meristem (RAM), leaves, cambium, embryos, and nodules in diverse angiosperm species.^23,24^ These analyses have revealed novel, cell type-specific gene expression patterns, even within rare cell types such as the quiescent center cells in RAM, leading to the discovery of crucial mechanisms underlying organ differentiation.^25,26^ In bryophytes, single-cell RNA-seq analysis in *Marchantia polymorpha* has revealed cell type-specific gene expression profiles along aging trajectories.^27^ However, the transcriptome of the SAM at the single-cell resolution in bryophytes is still unknown. In this study, we applied single-cell RNA-seq technology to *Physcomitrium patens* (*P. patens*), a model bryophyte suitable for stem cell studies, to uncover the regulatory mechanisms govering pluripotent stem cells and their origin.^28–30^

## Results

### Construction of a single-nucleus transcriptome atlas of gametophyte tissue in *Physcomitrium patens*

The gametophytic plant body of *P. patens* consists of filamentous tissues (protonemata) and leafy shoots (gametophores) (Figure 1A).^28,29^ Protonemata grow by tip growth of protonema apical cells and generation of protonemal cells backward through division of the apical cell. Protonemal cells frequently make side-branch initial cells. Most of the side branch initials become new protonemal apical cells, expanding the protonemal tissue area, while some are specified as a gametophore stem cell (gametophore apical cell).^4,29,30^ The gametophore apical cell undergoes asymmetric cell divisions with an oblique cell division plane.^4^ The apical daughter cell is maintained as a gametophore apical cell, while the basal daughter cell (merophyte) begins to differentiate. This oblique division of the gametophore apical cell continues cylindrically, forming a tetrahedral gametophore apical cell and creating the spiral topology of merophytes. The merophyte generates apical and basal daughter cells: the former forms a single leaf (phyllid) or an axillary hair, while the latter forms stem or rhizoid tissues. The gametophore apical cell, merophyte, and the derivatives of the merophyte are active in cell divisions, constituting the SAM.^10,15^ The gametophore apical cell remains at the tip of the SAM throughout the gametophore development. Unlike the unidirectional cell divisions of the protonema apical cell that produce only protonemal cells (2D growth), the gametophore apical cell exhibits pluripotent abilities, forming three-dimensional structures with various cell types (3D growth).^14,30^ This transition from a unipotent to a pluripotent state, along with its regular developmental patterns, makes the gametophore apical cell development an ideal model for studying mechanisms governing the specification of pluripotent stem cells in land plants.

**Figure 1.**
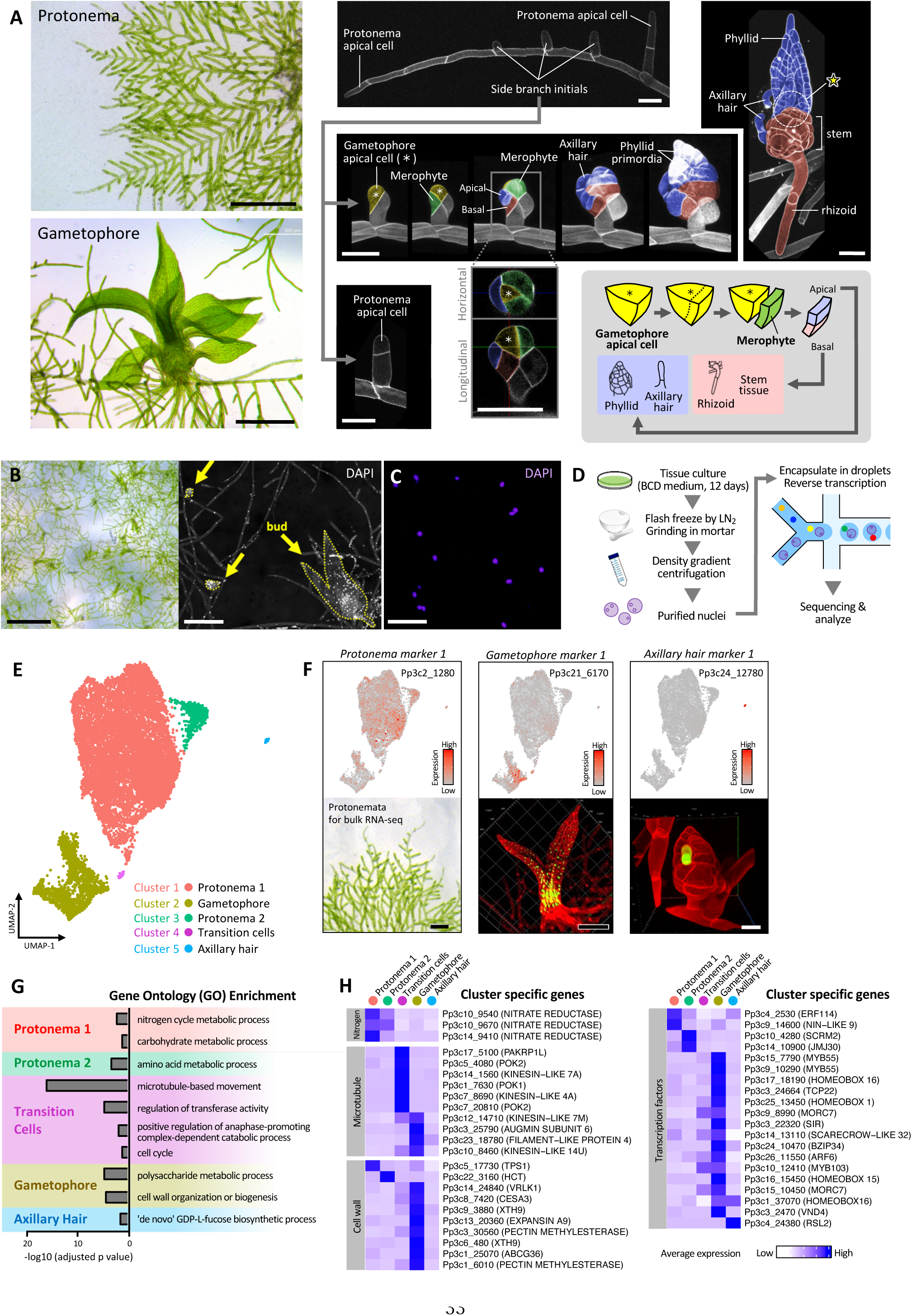
Construction of a single-nucleus transcriptome atlas of gametophyte tissue in *Physcomitrium patens*. (A) Developmental processes of *P. patens*. Bright field microscopy images of protonemata (upper) and a gametophore (bottom) are shown on the left, and confocal microscopy images on the right. Cell walls were stained with propidium iodide (PI) in the prior to confocal microscopy observation. The schematic diagram describes cell fate transition in the shoot apical meristem. The gametophore apical cell and a single cell-stage merophyte are shown in yellow and green, respectively. The apical and the basal daughter cells, shown as purple and blue, give rise to phyllid/axillary hair and stem tissues/rhizoid, respectively. The yellow star indicates the shoot apical meristem. Black scale bars, 500 µm. White scale bars, 50 µm. (B) Bright field (left) and the confocal microscopy images (right) of gametophyte tissue of *P. patens* used for single nuclei RNA-seq (snRNA-seq). Nuclei were stained with DAPI. Yellow arrows indicate young gametophore buds. Black scale bars, 1 mm. White scale bars, 200 µm. (C) Isolated nuclei stained with DAPI. Scale bar, 50 µm. (D) Workflow of snRNA-seq experiments. (E) Integrated uniform manifold approximation and projection (UMAP) plot displaying five cell clusters. Each dot represents a cell. (F) Expression of cell type-specific marker genes. Expression in the snRNA-seq data (top panels), plant tissues used for bulk RNA-seq (*Protonema marker 1*, Pp3c2_1280, left bottom panel), and the expression of promoter GFP lines for the other two markers (right bottom panels). The GFP signal is shown in green, and the cell wall is visualized by PI staining (red). Black scale bars, 200 µm, and a white scale bar, 20 µm. (G) Gene ontology (GO) terms (biological process) enriched in cluster-specific genes. (H) Average expression levels of cluster-specific genes related to nitrate metabolism, microtubules, cell wall biosynthesis, and transcriptional regulation. Genes with the lowest q-value are selected from each functional category.

To generate a single-cell atlas from *P. patens* gametophyte tissues, we collected nuclei from plant tissues containing protonemata and gametophores at various growth stages (Figures 1B to 1D). Isolated nuclei were processed using microfluidic device-based platforms and analyzed using a standard single-nucleus RNA-seq (snRNA-seq) experiment pipeline.^31^ We captured a total of 21,832 cells from the three independent replicates, capturing 1,005 genes per cell on average across the three replicates (Figure S1A). This performance is comparable to snRNA-seq results from other plant species.^32,33^ After filtering out low-quality cells, 14,454 cells were used for detailed analyses (Figure S1B).

The heterogeneity and homogeneity of filtered cells in the transcriptome were visualized using the uniform manifold approximation and projection (UMAP) and Leiden clustering. This analysis revealed five distinct cell clusters (Clusters 1-5) (Figure 1E). To identify the cell type of each cluster, we searched for genes with tissue-specific expression patterns in gametophytes as candidate markers from the reported transcriptome data.^13,22^ We then examined the expression patterns of these candidate marker genes by generating promoter-reporter lines. This approach led to the identification of four gametophore marker genes, two axillary hair marker genes, and one SAM marker gene (Table S1, Figures 1F and S1C). Additionally, we conducted bulk RNA-seq analyses using protonemata and young gametophores (Figures S2A and S2B). Genes highly expressed in protonemata found in the bulk RNA-seq data were used as protonema markers (Table S1). The largest cluster identified was Cluster 1, which comprises 11,925 cells, followed by Cluster 2, which had 1,821 cells. Given that the tissues analyzed were predominantly protonemata and young gametophores, we hypothesized that Cluster 1 and 2 correspond to the protonemal and gametophore cells respectively. To test this, we examined the expression patterns of cell type-specific marker genes (Table S1). As expected, the protonema marker gene was highly expressed in Clusters 1 and 3, while the gametophore marker gene showed strong expression in Cluster 2 (Figures 1F, S1C, S2C, and S2D). These findings suggest that Clusters 1 and 3 correspond to the protonemal cells and Cluster 2 to gametophore cells. Therefore, we assigned Clusters 1 and 3 as Protonema 1 and Protonema 2, respectively. The average expression levels of detected genes in the protonema and gametophore clusters were correlated well with bulk RNA-seq data, further validating these cell type assignments (Figure S2E). Cluster 4, positioned between the protonemal and gametophore cell clusters, likely represents cells transitioning from protonemata to gametophores (Figure 1E). Three gametophore markers (*Gametophore markers 1* through *3*) were also expressed in Cluster 4, and their promoter-reporter lines showed expression from a very early stage (two or four-cell stage) of gametophore development (Figure S1C). In contrast, we hardly detected expression of the promoter-reporter line of *Gametophore marker 4* in the two-cell stage gametophore. The *Gametophore marker 4* showed weaker expression in Cluster 4 (Figure S1C). These observations further support that Cluster 4 represents cells transitioning from protonemata to gametophores. Additionally, the expression of axillary hair marker genes was enriched in Cluster 5, a small, distinct cluster, indicating that Cluster 5 corresponds to axillary hair cells (Figures 1F and S1C). Overall, the snRNA-seq analysis efficiently captured both transcriptomic heterogeneity and homogeneity among the major cell types in gametophyte tissues in *P. patens*.

We analyzed the transcriptomic characteristics of each cluster by identifying cluster-specific genes. We detected 67 to 449 specific genes in each cluster (Table S1). Gene ontology (GO) enrichment analysis revealed significant enrichment of biological functions associated with each set of cluster-specific genes (Figure 1G). Specifically, the protonema 1 cluster exhibited enrichment of GO terms ‘nitrogen cycle metabolic process’, which correlates with the significant role of nitrate in protonemal growth (Figure 1G).^34^ This enrichment includes up-regulation of orthologs of *NITRATE REDUCTASE*, a gene coding for an enzyme required for nitrate assimilation (Figure 1H).^35^ In the transition-cell cluster, GO terms related to microtubules and/or cell cycle were significantly enriched (Figure 1G). Notably, *PHRAGMOPLAST ORIENTING KINESIN* (*POK*), a key factor in determining the orientation of the cell division plane, was specifically up-regulated (Figure 1H).^36^ The up-regulation of *POK* in cells transitioning from protonemal to gametophore cells suggests its importance in side branch initial formation from protonemal cells and the first oblique cell division of the gametophore apical cell.^37^ In the gametophore cluster, cluster-specific genes were enriched in cell wall-related GO terms (Figure 1G). Specifically, orthologs of *CELLULOSE SYNTHASE A* (*CESA*), *XYLOGLUCAN ENDOTRANSGLUCOSYLASE/HYDROLASE* (*XTH*), and *PECTIN METHYLESTERASE* were significantly up-regulated in the gametophore cluster (Figure 1H). The up-regulation of CESA orthologs underscored their essential role in gametophore development.^38^

We also explored the transcription factors included in the cluster-specific genes (Figure 1H). In the protonema 1 cluster, an ortholog of *NIN-LIKE PROTEIN*, closely related to the nitrate sensor, was up-regulated, consistent with the increased expression levels of nitrate metabolic enzymes.^39^ Conversely, many orthologs of transcription factors related to development in angiosperms, such as *HOMEOBOX1* (*HB1*), *HB15*, *HB16*, *MYB103*, and *VASCULAR RELATED NAC-DOMAIN 4* (*VND4*), were up-regulated in the gametophore cluster.^40–45^ The increased expression levels of transcription factors involved in plant development in angiosperms in the gametophore cluster likely reflect the transition from filamentous protonema development to complex leafy shoot development. In the axillary hair cluster, an ortholog of *RHD SIX-LIKE* (*RSL*), involved in root hair development in angiosperms, was specifically up-regulated.^46^

### Dynamic changes in gene expression during SAM development

The *P. patens* SAM localizes in the gametophore and is covered by phyllids (Figure 1A). We focused on the cells within the gametophore cluster to further dissect the transcriptome profiles during SAM development. The meristem marker gene we identified (Pp3c27_870) showed clear expression in the meristem region of the gametophore from the first division of the gametophore apical cell, and the expression continued in the SAM region in the gametophore shoots (Figure 2A). The meristem marker gene was expressed at the right corner of the gametophore cluster in the UMAP plot (Figure 2B). Conversely, an ortholog of *Class III Homeodomain-Leucine Zipper* (*PpC3HDZ1*), expressed explicitly in the phyllid primordia, was highly detected in cells in the lower region of the gametophore cluster.^47^ An ortholog of *Class I Homeodomain-Leucine Zipper* (*Pphb7*), known to be expressed in stem tissues including rhizoid, was expressed in cells in the upper part of the gametophore cluster (Figure 2B).^48^ Other genes known to be expressed in phyllid primordia or stem tissues also showed expression patterns similar to *PpC3HDZ1* and *Pphb7* in the UMAP plot, respectively (Figures S3A and S3B).^5,47^ These results indicate that the undifferentiated meristematic cells are located around the right corner of the gametophore cluster, cells differentiating into phyllids are located in the lower part, and cells differentiating into stem or rhizoid (stem/rhizoid) are located in the upper part of the gametophore cluster. Among the four gametophore markers, three (Pp3c21_6170, Pp3c12_13360, Pp3c7_3350) were expressed in the gametophore from the two or four-cell stage, while the expression of *Gametophore marker 4* (Pp3c3_34230) started from the gametophore at the several-cell stage (Figure S1C). Consistently, the localization of cells expressing Pp3c3_34230 showed a slight shift compared to that of the other three markers. Additionally, an ortholog of *LONELY GUY* (*PpLOG1*), a gene known to have gametophore apical cell-specific expression, was specifically expressed at the right corner of the meristematic region in the gametophore cluster (Figure 2C).^49^ These findings support our notion that the gametophore apical cells were localized at the right corner of the gametophore cluster.

**Figure 2.**
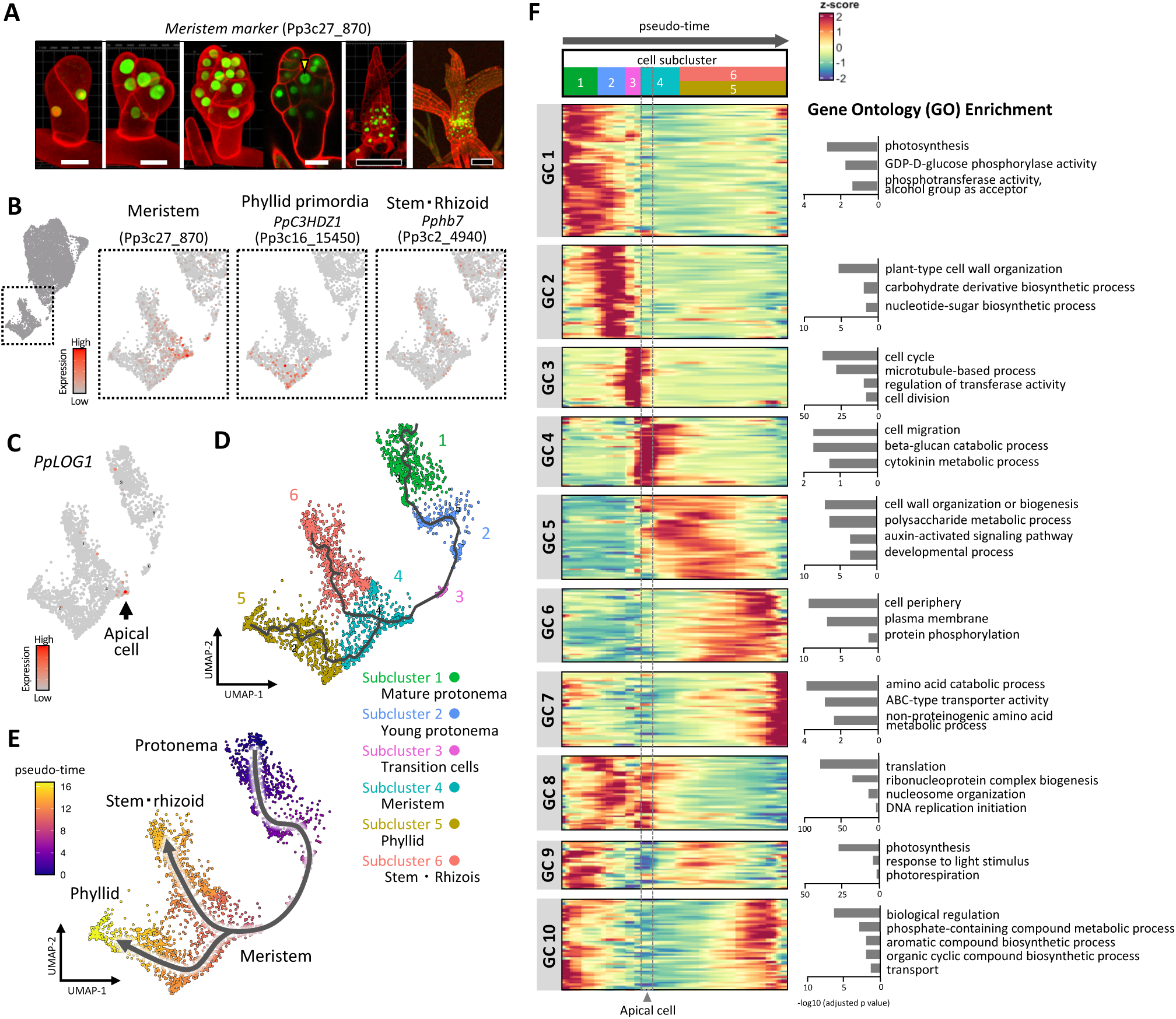
Dynamic changes in gene expression during SAM development. (A) Expression patterns of the meristem marker gene (Pp3c27_870) during gametophore development. The GFP signal driven by the promoter of the marker gene is green, and the cell wall visualized by PI staining is red. Yellow arrowhead: gametophore apical cell. White scale bars, 20 µm. Black scale bars, 100 µm. (B) Expression patterns of *Meristem marker* (Pp3c27870), *PpC3HD-ZIP1* (Pp3c16_15450), and *Pphb7* (Pp3c2_4940) in the gametophore cell cluster (indicated by dashed-line rectangle) of UMAP plot. *PpC3HD-ZIP1* and *Pphb7* are known to be expressed in phyllid primordia and stems/rhizoids, respectively.^47,48^ Cells around the gametophore cell cluster (dashed-lined rectangle) are magnified. (C) The expression pattern of *PpLOG1*, which localizes to the gametophore apical cells (black arrow) around the gametophore cluster in the UMAP plot. (D) Extracted cells around the trajectory from protonema cells to gametophyte cells. Black solid lines indicate cell fate trajectory predicted from gene expression dynamics. (E) Pseudotime calculated from the protonema to gametophore trajectory. The initial point of the pseudotime is set at the terminus of trajectory in the protonema cells. (F) Left: Expression patterns of genes that show significant changes (q value < 0.05) along pseudotime. Z-scores of expression levels during the pseudotime were calculated for the detected genes. The detected genes were classified into ten gene clusters (GC1 to GC10) by k-means clustering (k = 10). The horizontal axis is pseudotime (from protonema to gametophore, left to right). Numbers at the top of the heatmap correspond to the UMAP plot’s subcluster number shown in C. Dashed gray lines indicate gametophore apical cells enriched pseudotime, the earliest region of the pseudotime in Subcluster 4. Right: Significant GO terms enriched in each GC. Only ‘driving terms’ detected by g:profiler software are shown to avoid redundancy.

The phyllid primordia and stem/rhizoid are produced from the merophyte, which derives from the daughter cell of the asymmetric division of the gametophore apical cell (Figure 1A). The phyllid and the stem tissue cell lineage are separated after the first division of the merophyte. Therefore, we next tested if the gametophore cells in snRNA-seq data follow these cell lineages. We inferred cell fate transition using trajectory detection based on the transcriptomic changes of each cell. A trajectory map was constructed connecting the protonema, transition cells, and gametophore clusters (Figures 2D and S3C). The trajectory passed through the meristematic region in the gametophore cluster and immediately branched into the phyllid and stem/rhizoid tissue cell populations. The predicted trajectory reflected the known developmental patterns of SAM.

To examine the changes in gene expression during the cell fate transition around the gametophore apical cell in detail, we extracted cells located around the trajectory from the protonema to the gametophore on the UMAP plot using Monocle3 software. The extracted cells were divided into six subclusters (Subcluster 1-6), using Leiden clustering (Figure 2D). Subclusters 1 and 2 are associated with protonemal cells. Subcluster 2, positioned closely to the transition cells, likely contains young protonemal cells capable of forming side branch initials, whereas Subcluster 1 contains mature protonemal cells. Subcluster 3 corresponded to the transition cell cluster (Figures 1E and 2D). Subcluster 4, 5 and 6 represents meristematic tissues of the gametophore, phyllid, and stem/rhizoid tissues respectively (Figures 2B and 2D). We performed pseudotime analyses by setting the initial position of pseudotime at the terminus of the trajectory located in the protonemal cells. Each cell was assigned a pseudotime value relative to the initial position of the trajectory (Figure 2E). Continuous progression of the pseudotime suggests the dynamic changes of gene expression profiles from protonemata to gametophores, including the gametophore apical cell development.

We identified 5,785 genes showing significant change in expression along pseudotime (Table S1). These genes were classified into ten gene clusters (GC1 - GC10) based on their expression patterns along pseudotime (Figure 2F). Genes in GC1 to GC7 were up-regulated at specific developmental stages during cell fate transition, from protonema (Subcluster 1 and 2) to gametophore (Subcluster 4, 5, and 6) through transition and bud initiation (Subcluster 3). GC1 genes showed high expression in Subcluster 1, corresponding to mature protonemal cells (Figure 2F). Enrichment of GO terms related to photosynthesis is consistent with the characteristics of mature protonemal cells, which possessed well-developed chloroplasts. GC2 genes were up-regulated in cells of Subcluster 2, likely representing young protonemal cells (Figure 2F). GC2 showed enrichment of cell wall-related GO terms. Orthologs of *EXPANSIN A*, which promotes side branch initial formation in protonemata, and an ortholog of *COBLA LIKE PROTEIN 7*, involved in determining tip growth polarity, were up-regulated (Table S1).^17,50^ GC3 genes were up-regulated during the transition stage (Figure 2F). GO terms related to microtubules, cell cycle, and cell divisions were highly enriched. Notably, *targeting protein for Xklp2* (*TPX2*), a critical regulator of the first oblique cell division of gametophore apical cell, was included in GC3 (Table S1).^51^ GC4 genes were activated during gametophore bud initiation and meristem formation (Figure 2F). In GC4, the GO term ‘cytokinin metabolic process’ was significantly enriched, and orthologs of *LOG*, encoding an enzyme catalyzing the final steps of active cytokinin production, were included (Table S1).^49^ Genes in GC5 were activated immediately after GC4 in the pseudotime during merophyte development (Figure 2F). Enrichment of GO terms such as ‘cell wall organization or biogenesis’, ‘auxin-activated signaling pathway’, and ‘developmental process’ were detected. *PpSHORTROOT (PpSHR)*, encoding the GRAS transcription factor expressed in phyllid primordia and regulating midrib formation, was up-regulated (Table S1).^52^ Furthermore, GC5 included orthologs of *EPIDERMAL PATTERNING FACTOR LIKE* (*EPFL*) and *HB1*, important regulators of shoot development in angiosperms.^40,53^ GC6 genes were activated later than GC5 in pseudotime (Figure 2F). A GO term, ‘cell periphery’ was significantly enriched. Transporters involved in cell wall biosynthesis, such as *Exocyst complex component EXO70A1*, and genes involved in secondary cell wall formation in angiosperms, such as *VASCULAR-RELATED RECEPTOR-LIKE KINASE* (*VRLK*) and *WALLS ARE THIN* (*WAT*) were up-regulated (Table S1).^54–56^ These results suggest further progression of cell differentiation and completion of organogenesis. GO terms related to molecule transport were enriched in GC7 genes, which are activated at the end of pseudotime (Figure 2F). Orthologs of transporters involved in stress response or tolerance, such as *PLEIOTROPIC DRUG RESISTANCE 11* (*PDR11*) and *ABC transporter C family 5* (*ABCC5*), were also up-regulated in GC7 genes, suggesting the establishment of responses to environmental stress after morphogenesis (Table S1).^57,58^ GC8, 9, and 10 genes were activated during protonemal and gametophore growth at two developmental timings (Figure 2F).

### Genes associated with auxin and morphogenesis are up-regulated during merophyte development

We demonstrated that Subclusters 5 and 6 represent differentiated tissues such as phyllids and stems/rhizoids derived from merophytes. In contrast, Subcluster 4 corresponds to the SAM, including the gametophore apical cell (Figures 2C and 2D). To further elucidate the differences between pluripotent gametophore apical cells and differentiating merophytes, we investigated the types of genes activated during merophyte development. The GC5 genes, likely indicative of early merophyte development, included auxin-related genes such as *AUXIN RESPONSE FACTOR* (*ARF*) and *PINFORMED* (*PIN*) (Figure 2F and Table S1).^19,59,60^ We identified auxin biosynthesis and signaling genes among those showing significant expression changes during pseudotime (Figure 3A). Orthologs of *AUXIN SIGNALING F-BOX 1* (*PpAFB1*) and *PpAFB2*, which are auxin receptors, were up-regulated throughout the gametophore cluster (Figures 3A and 3B).^61^ Conversely, most auxin signaling and biosynthesis genes, including *AUXIN/INDOLE-3-ACETIC ACID* (*PpIAA*s), *ARF* (*PpARF*s), *TRYPTOPHAN AMINOTRANSFERASE RELATED* (*PpTAR*s), *YUCCA* (*PpYUC*), *PIN* (*PpPIN*s), *AUXIN TRANSPORTER PROTEIN* (*PpAUX1/LAX*s), and *GRETCHEN HAGEN 3* (*PpGH3*s) showed up-regulation later than the apical cell stage during pseudotime (Figures 3A and 3B).^19,59–65^ These findings suggest that the entire gametophore region was capable of auxin perception; components involved in auxin synthesis and signaling were activated explicitly during merophyte development (Figure 3C).

**Figure 3.**
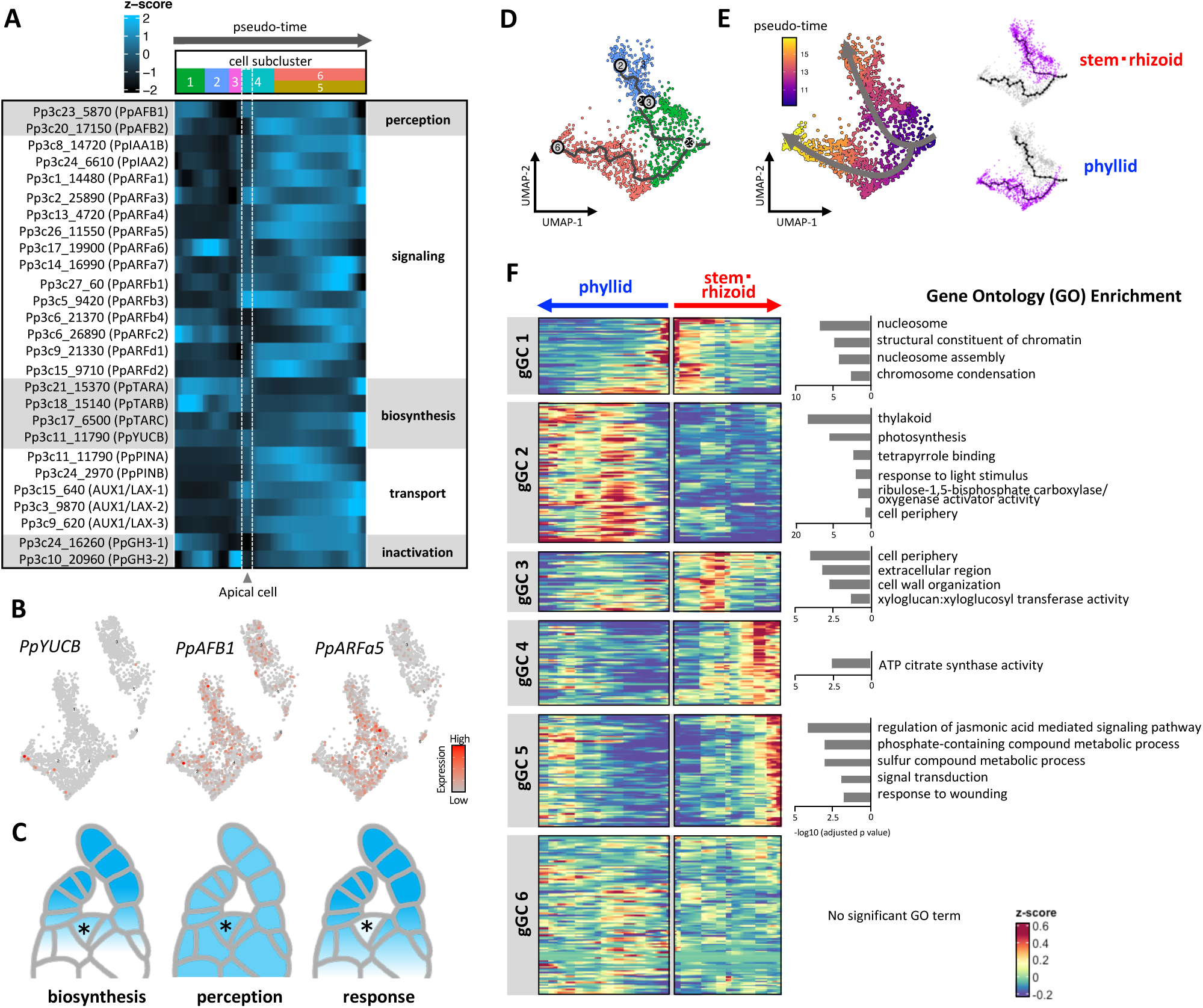
Genes associated with auxin and morphogenesis are up-regulated during merophyte development. (A) Expression patterns (z-score) of auxin biosynthesis and signaling genes during pseudotime (horizontal axis). Dashed white lines indicate gametophore apical cells enriched pseudotime, the earliest region of the pseudotime in Subcluster 4. (B) Expression patterns of representative genes involved in auxin biosynthesis (*PpYUCB*), perception (*PpAFB1*), and signaling (*PpARFa5*) in the UMAP plot. (C) Schematic summary of the spatial distribution of auxin biosynthesis, perception, and response (shown with blue color) in the *P. patens* SAM as deduced from data presented in Panel B. The asterisk indicates the gametophore apical cell. (D and E) UMAP plot of cells in the trajectory from the initiation at the branching point to the terminus in the gametophore cluster. The trajectory (solid black lines in D) and the pseudotime transition after the trajectory branching (E) are shown. Cells in the stem/rhizoid or phyllid trajectory are indicated magenta. (F) Left: Expression patterns (z-score) of genes differentially expressed during pseudotime from the branching point to the terminus (q value < 0.01). Identified genes were classified into six gametophore gene clusters (gGC1 to gGC6) by k-means clustering (k = 6). The horizontal axis is the pseudotime (from the middle to the right in the stem/rhizoid trajectory; from the middle to the left in the phyllid trajectory). Right: Significant GO terms enriched in each gGC. Only ‘driving terms’ detected by g:profiler software are shown to avoid redundancy.

After the stage involving the apical cell, the developmental trajectory bifurcates into phyllids and stems/rhizoids (Figure 2D). To further elucidate merophyte development, we categorized gametophore cells into the phyllid and stem/rhizoid tissue trajectories (Figure 3D). In our analysis, we defined the branching point of these trajectories as the initiation of pseudotime (Figure 3E). We identified 2,208 genes exhibiting significant expression changes during the pseudotime (Table S1). These genes were classified into six gametophore gene clusters (gGC1 - gGC6) based on their expression patterns in the pseudotime (Figure 3F). Specifically, gGC1 genes were up-regulated at the initial pseudotime point of the trajectory. Notably, GO terms related to nucleosome and chromatin were significantly enriched (Figure 3F). Orthologs of *CHROMATIN REMODELING 5* (*CHR5*) and *CHROMOMETYLASE 2* (*CMT2*), involved in chromatin condensation and gene silencing in angiosperms, were up-regulated (Table S1).^66,67^ gGC2 genes were exclusively up-regulated in the phyllid trajectory post-branching point (Figure 3F). Consistently, GO terms ‘thylakoid’ and ‘photosynthesis’, likely related to chloroplast maturation, were significantly enriched. Additionally, several orthologs of genes known as key regulators of shoot development in angiosperms, such as *ERECTA LIKE 1* (*ERL1*), *LATERAL BOUNDARY DOMAIN 12* (*LBD12*), and *HB33* were up-regulated (Table S1).^53,68,69^ In the stem/rhizoid trajectory, gGC3 genes initially showed up-regulation post-branching point (Figure 3F). gGC3 genes displayed enrichment of GO terms ‘cell periphery’ and ‘cell wall organization’, suggesting the importance of cell wall biosynthesis or signaling at the plasma membrane. The gGC3 genes include an ortholog of *CESA*, a cell wall biosynthesis gene required for gametophore development, and an ortholog of *CLAVATA1* (*PpCLV1a*), which represses gametophore apical cell formation on the stem (Table S1).^5,38^ Following gGC3, gGC4 genes were up-regulated (Figure 3F). The gGC4 genes include orthologs of *HB1* and *HB16*, known as regulators of leaf development, but their function in *P. patens* remains unknown (Table S1).^40,43^ gGC5 genes showed increased expression at the terminus of the stem tissue trajectory (Figure 3F). GO terms ‘regulation of jasmonic acid mediated signaling pathway’ and ‘response to wounding’ were enriched in this gene cluster, indicating increased expression of genes related to environmental stress.

### Cytokinin biosynthesis is specifically localized to the apical cell, while perception occurs ubiquitously in the gametophore

It is well-established that cytokinins promote gametophore apical cell formation of *P. patens*.^20,70^ To further investigate the localization of cytokinin signaling during the cell fate transition around the gametophore apical cell, we identified genes involved in cytokinin biosynthesis and perception among genes showing significant changes during pseudotime (Figure 4A). Orthologs of *CHASE DOMAIN HISTIDINE KINASE* in *P. patens* (*PpCHK1, PpCHK2, PpCHK3*), which are cytokinin receptors, were up-regulated throughout the gametophore cluster (Figures 4A and 4B).^71^ Conversely, orthologs of *LOG* (*PpLOG1 - PpLOG3*), which are enzymes of active cytokinin biosynthesis, were specifically up-regulated in the gametophore apical cell position (Figure 4A and 4B).^49^ Further examination of *PpLOG2* promoter activity confirmed a strong localization to the gametophore apical cell similar to that of *PpLOG1* (Figure 4C).^49^ Given that *PpCHK1, PpCHK2,* and *PpCHK3* were uniformly expressed in the gametophore cluster, cytokinin response localization to the gametophore apical cell likely depends on PpLOG localization (Figure 4D).^49^ The snRNA-seq data also revealed up-regulation of two *CYTOKININ OXIDASE* genes in *P. patens* (*PpCKX4* and *PpCKX5*) at the apical cell position in the pseudotime, suggesting strict regulation of cytokinin levels at the gametophore apical cell (Figure 4A).^72^ Additionally, two orthologs of *ISOPENTENYL TRANSFERASE* (*PpIPT3* and *PpIPT5*) were expressed in the stem region of the gametophore cluster (Figures 4A and 4B).^73^ Cytokinin biosynthesis regions in the gametophore tissue might be compartmentalized based on reaction steps (Figure 4D).

**Figure 4.**
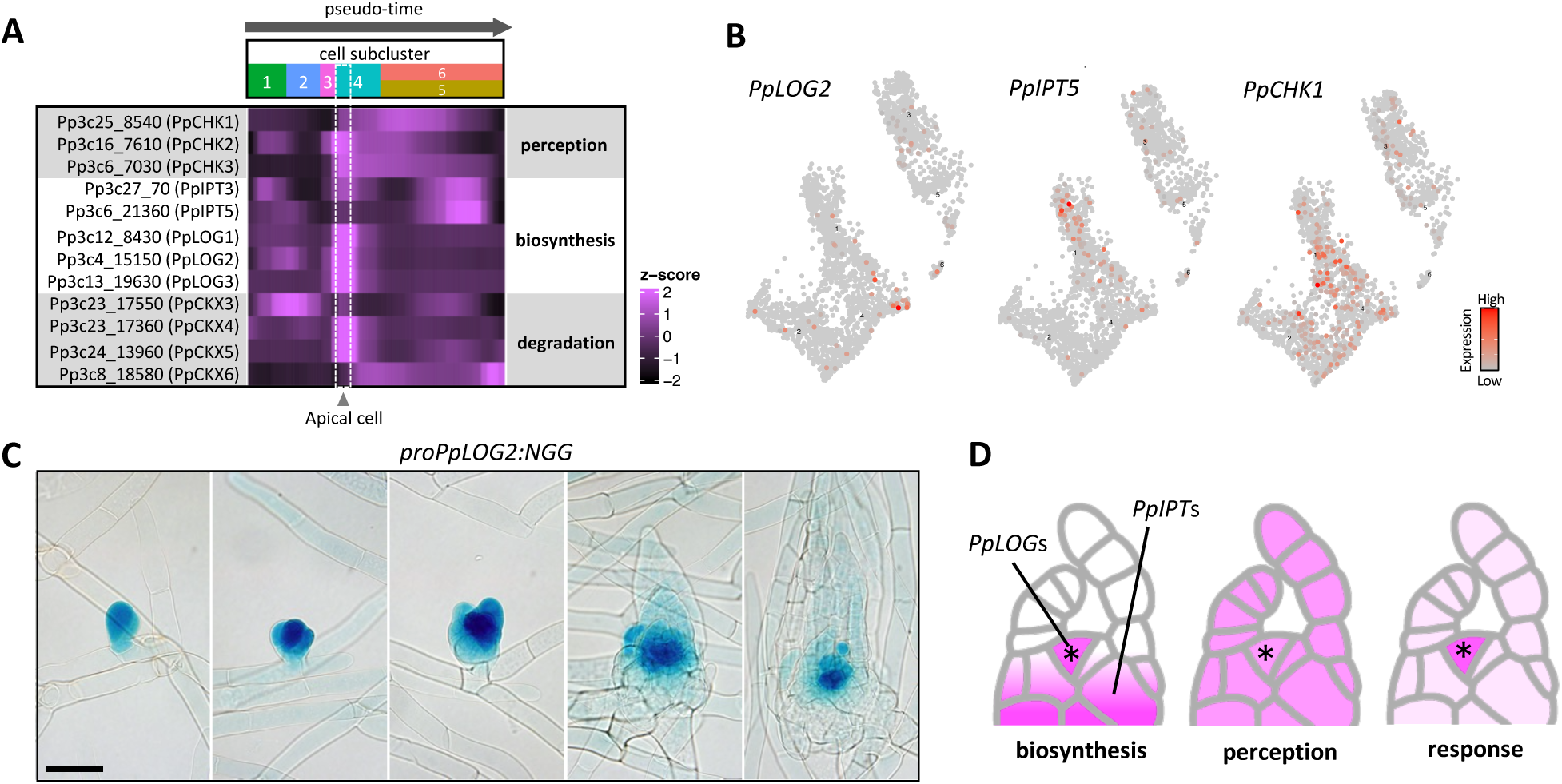
Cytokinin biosynthesis is specifically localized to the apical cell, while perception occurs ubiquitously in the gametophore. (A) Expression patterns (z-score) of cytokinin biosynthesis and signaling-related genes differentially expressed during pseudotime during pseudotime from the branching point to the terminus (q value < 0.01). The horizontal axis of the heatmap represents the pseudotime. Dashed white lines indicate gametophore apical cells enriched pseudotime, the earliest region of the pseudotime in Subcluster 4. (B) Expression patterns of genes representative of cytokinin biosynthesis (*PpLOG2* and *PpIPT5*), and signaling (*PpCHK1*) in the UMAP plot. (C) Expression patterns of *PpLOG2* promoter-NLS-GFP-GUS (NGG) reporter gene. Scale bar, 50 µm. (D) Schematic summary of the spatial distribution of cytokinin biosynthesis, perception, and response (shown with magenta color) in *P. patens* SAM as deduced from data shown in Panel B. The asterisk indicates the gametophore apical

### The cytokinin-ESR module promotes the formation of gametophore apical cells

We then focused on identifying transcription factors significantly up-regulated in the gametophore apical cell to pinpoint key regulatory genes essential for its initiation. Among the 5,785 genes showing significant expression changes from protonema to gametophore pseudotime, 325 were transcription factors. Based on their expression patterns during pseudotime, these transcription factors were classified into ten groups (Figure S4A). We found that two out of the ten groups (Groups 3 and 9), comprising 38 transcription factors in total, exhibited specific up-regulation around the gametophore apical cell position (Figure S4A). These 38 transcription factors were further sorted by k-means clustering based on expression patterns, and 28 transcription factors that exhibit specific activation at the gametophore apical cell were identified (Figure 5A). These apical cell-specific transcription factors included orthologs of *APETALA PLETHORA BABYBOOM* (*APB2* and *APB3*), crucial for the gametophore apical cell formation (Figure 5A).^74^ While their functions in *P. patens* remain unknown, orthologs of *BES1/BZR1 HOMOLOG PROTEIN 1* (*BEH1*) involved in brassinosteroid signaling, as well as several *AP2/ERF* family and *MYB* family transcription factors were also identified. To further elucidate the key factors govering gametophore apical cell identity, we focused on downstream genes influenced by cytokinins, which are known to promote gametophore apical cell formation.^20,70^ Initially, we identified cytokinin-responsive genes using bulk RNA-seq analyses of plant colonies treated with 6-benzylaminopurine (BAP), a synthetic cytokinin (Figure S4B). A total of 336 genes were identified as differentially expressed genes (DEGs), with the majority being up-regulated upon cytokinin application (Figure S4C and Table S1). Cytokinin responsiveness of gametophore apical cell-specific transcription factors is shown in the Figure 5A. Notably, the ortholog of *ENHANCER OF SHOOT REGENERATION1/DORNROSCHEN* (*PpESR*s), an *AP2/ERF* family transcription factor, showed the most remarkable up-regulation in response to cytokinin among 28 gametophore apical cell-specific transcription factors (Figure 5A).

**Figure 5.**
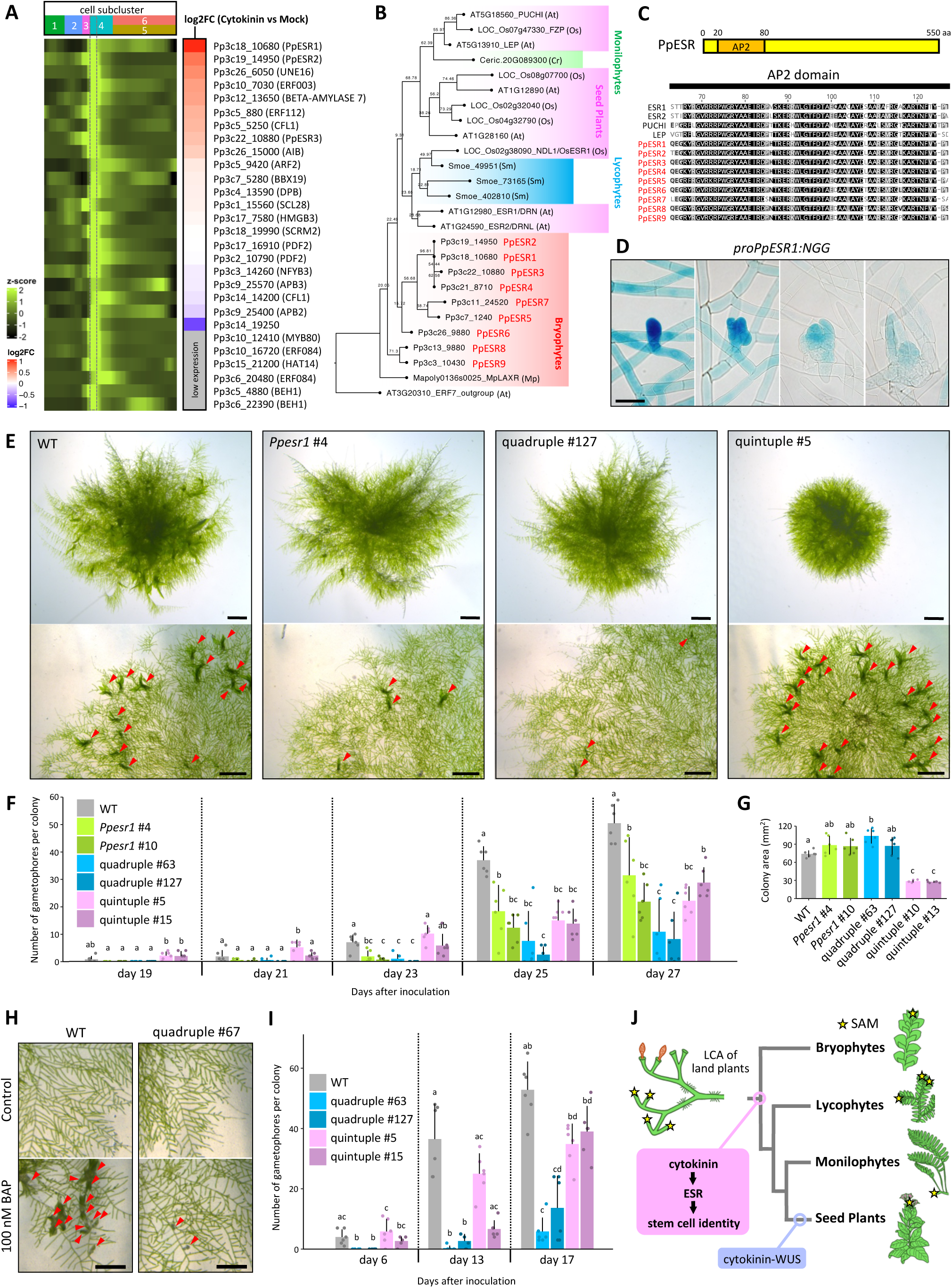
The cytokinin-ESR module promotes the formation of gametophore apical cells. (A) Expression patterns (z-score) of transcription factors up-regulated in the gametophore apical cell. The horizontal axis of the heatmap represents the pseudotime. Dashed gray lines indicate gametophore apical cells enriched pseudotime. The color code from blue to red indicates gene expression changes (Log2 fold change value) after cytokinin (BAP) treatment as revealed by bulk RNA-seq. Genes with an M value of less than 0.1 in bulk RNA-seq are considered low-expression genes. Genes from the AP2/ERF family are indicated by red asterisk. (B) Phylogenetic tree of the ESR1/DRN clade in subfamily VIII, which belongs to the AP2/ERF family of transcription factors.^109^ Genes are from *Physcomitrium patens* (PpESR1-PpESR9), *Marchantia polymorpha* (Mp), *Selaginella moellendorffii* (Sm), *Ceratopteris richardii* (Cr), *Arabidopsis thaliana* (At), and *Oryza sativa* (Os). ERF7, which falls outside of the ESR1/DRN clade, was included as an outgroup. The bootstrap values are indicated at each branch point. (C) Domain structure of PpESR proteins and an alignment of amino acid sequences of AP2 domain from PpESRs and AP2/ERF proteins related to ESR1/DRN from *A thaliana*. (D) Expression patterns of *PpESR1* promoter-NLS-GFP-GUS (NGG) reporter gene. Scale bar, 50 µm. (E) Growth phenotypes of *Ppesr1* single mutants, *Ppesr1 Ppesr2 Ppesr3 Ppesr4* quadruple mutants, and *Ppesr5 Ppesr6 Ppesr7 Ppesr8 Ppesr9* quintuple mutants. Upper panels are plants grown on BCDAT medium for 13 days, and lower panels are plants grown between two cellophane membranes on BCD medium for 27 days. Gametophores are indicated by red arrowheads. Scale bars, 1 mm. (F) Number of gametophores per colony of the wild type (WT) and *Ppesr* mutants grown between two cellophane membranes on BCD medium. Statistical significance was evaluated by by HSD test (n = 6, p < 0.05). (G) Colony area of the WT and *Ppesr* mutants grown between two cellophane membranes on BCD medium for 21 days. Statistical significance was evaluated by HSD test (n = 6, p < 0.05). (H) Growth phenotype of WT and *Ppesr1 Ppesr2 Ppesr3 Ppesr4* quadruple quadruple mutant grown between two cellophane membranes on BCD medium supplemented with cytokinin (BAP, 100 nM) for 13 days. Gametophores are indicated by red arrowheads. Scale bars, 500 µm. (I) Number of gametophores per colony of the wild type (WT) and *Ppesr* mutants grown between two cellophan membranes on BCD medium supplemented with cytokinin (BAP, 100 nM). Statistical significance was evaluated by HSD test (n = 6, p < 0.05). (J) Evolution of molecular pathways underlying the pluripotent stem cell identity in land plants. The cytokinin-ESR pathway is conserved between bryophytes and angiosperms, suggesting that this pathway is an ancient module that originated in the common ancestor of land plants. Other stem cell factors known in angiosperms, such as WUS, were recruited to stem cell regulation after the diversification of vascular plants. The illustration of the last common ancestor (LCA) of land plants is based on Kenric 2018 with some modifications.^6^

*ESR*s are highly conserved in land plants (Figure 5B). They function downstream of auxin and cytokinin, promoting stem cell formation in the SAM during shoot regeneration in *A. thaliana*.^75–77^ *NARROW AND DWARF LEAF 1* (*NDL1*), an ortholog of *ESR* in rice (*Oryza sativa*), is essential for maintaining stem cells in the SAM.^78^ These findings led us to hypothesize that *ESR* may be an evolutionary conserved factor promoting pluripotent stem cell identity. We tested whether *ESR* is involved in promoting gametophore apical cell identity influenced by cytokinin in *P. patens*. The *P. patens* genome contains nine orthologs of *ESRs* (*PpESR1-PpESR9*) (Figure 5B). The amino acid sequences of the AP2 domain, crucial for DNA binding, are well conserved in the nine PpESRs (Figure 5C). Phylogenetic analysis indicates that *PpESR1 - PpESR4* form a distinct clade separate from *PpESR5 - PpESR9* (Figure 5B). The *PpESR1-PpESR4* genes contain a single intron, whereas *PpESR5 - PpESR9* have two introns (Figure S4D). Notably, only *PpESR1, PpESR2* and *PpESR3* were found among the 28 apical cell-specific transcription factors (Figure 5A). Thus, we hypothesized that *PpESR1 - PpESR4* primarily regulate gametophore apical cell development. The expression patterns of *PpESR1* were validated *in planta* using *proPpESR1:NLS-GFP-GUS* (*proPpESR1:NGG*) reporter lines, revealing the strongest GUS activity at the initiation stage of the gametophore apical cell, consistent with the snRNA-seq data (Figure 5D).

To investigate the role of *PpESR1-PpESR4* in gametophore apical cell development, we generated loss-of-function mutants using CRISPR/Cas9. We obtained independent two lines of *Ppesr1* single mutants, two lines of *Ppesr1 Ppesr2 Ppesr3 Ppesr4* quadruple mutants, and two lines of *Ppesr5 Ppesr6 Ppesr7 Ppesr8 Ppesr9* quintuple mutants (Figure S4E). The *ppesr1* mutants showed a significant delay in gametophore formation (Figures 5E and 5F). Gametophores formed in the *Ppesr1 Ppesr2 Ppesr3 Ppesr4* quadruple mutants, albeit with a more delay compared to the *Ppesr1* single mutants (Figures 5E and 5F), indicating that *PpESR1-PpESR4* are involved in promoting gametophore apical cell formation. In contrast, the *Ppesr5 Ppesr6 Ppesr7 Ppesr8 Ppesr9* quintuple mutants did not affect gametophore formation (Figures 5E and 5F). However, the colony size of the *Ppesr5 Ppesr6 Ppesr7 Ppesr8 Ppesr9* quintuple mutants was reduced compared to that of the wild type and to the *Ppesr1 Ppesr2 Ppesr3 Ppesr4* quadruple mutants (Figures 5E and 5G). The frequency of cell divisions in protonemal apical cell was significantly reduced in the *Ppesr5 Ppesr6 Ppesr7 Ppesr8 Ppesr9* quintuple mutants (Figure S5). These data suggest that *PpESR5-PpESR9* are involved in protonemal growth but not in the development of gametophore apical cells.

The up-regulation of *PpESR1* expression by cytokinin prompted us to investigate whether the role of cytokinins in promoting gametophore apical cell identity depends on the function of *PpESR1-PpESR4*. In both the wild type and *Ppesr5 Ppesr6 Ppesr7 Ppesr8 Ppesr9* quintuple mutants, gametophore formation was observed on day 6 after the addition of BAP. In contrast, bud formation was observed at day 13 in the *Ppesr1 Ppesr2 Ppesr3 Ppesr4* quadruple mutants grown under the same condition, indicating that *PpESR1-PpESR4* act as downstream regulators of cytokinin to promote gametophore apical cell initiation (Figures 5H and 5I).

## Discussion

The apical meristem, containing pluripotent stem cells, enables plants to continuously differentiate new organs throughout their lives.^9^ Originating in the common ancestor of land plants, the apical meristem has been a key innovation that allowed plants to flourish. ^2,6–8^ However, the fundamental nature of pluripotent stem cells in plants remains ambiguous. Taking advantage of *P. patens*, which features a single stem cell (the apical cell) within the SAM and is suitable for molecular genetic analysis including single cell imaging, we conducted single-cell RNA-seq analysis to elucidate the minimal and ancestral mechanisms underlying pluripotency. Our analysis identified a cell cluster at the protonema to gametophore transition, and a subcluster containing SAM in the gametophore cluster. Enriched GO terms in transition cells included cell cycle and cell division, contrasting with cytokinin metabolic process enrichment in apical cells. Cytokinins promote gametophore stem cell identity in *P. patens*.^20,70^ Cell divisions of the gametophore apical cell that give rise to the SAM is also triggered by cytokinin. We previously demonstrated that PpLOG1 specifically localizes in the apical cell, indicating that cytokinin biosynthesis is confined to the apical cell.^49^ Consistently, the localization of cytokinin in apical cell was demonstrated by using *TCSv2* as a molecular marker.^49^ In this study, we showed a more comprehensive distribution of gene expression involved in cytokinin biosynthesis and signaling. Not only *PpLOG1* but also several other paralogs of *PpLOG*s showed specific up-regulation in the gametophore apical cell. On the other hand, cytokinin receptors were ubiquitously expressed in the gametophore including the apical cell. Cytokinin degradation genes were specifically up-regulated in the apical cell, probably due to the feedback control, reflecting high cytokinin levels in the apical cell.^20^ These findings imply that the cytokinin signal is activated by apical cell-specific biosynthesis by PpLOGs, and simultaneously confined to the apical cell through the immediate degradation by PpCKXs. We identified four ESR1 orthologs (*PpESR1* to *PpESR4*) as downstream regulators of cytokinin, promoting gametophore apical cell identity. In *A. thaliana*, *ESR1* operates downstream of cytokinin during SAM regeneration from calli, suggesting that the cytokinin-ESR pathway may be an ancient mechanism promoting pluripotent stem cell identity that evolved in the common ancestor of land plants (Figure 5J).^75,76^ In *M. polymorpha*, *LOW-AUXIN RESPONSIVE* (Mp*LAXR*), an ortholog of *ESR1*, promotes shoot regeneration, although its relationship with cytokinin remains unclear.^79^

In contrast to cytokinin accumulation in the gametophore apical cell, genes involved in auxin signaling were up-regulated in the differentiating merophytes. Auxin promotes cell differentiation by inhibiting cell divisions but enhancing cell elongation and rhizoid formation in *P. patens*.^19,21,59^ Its levels are minimal in the gametophore apical cell but high in the merophyte.^80^ Our snRNA-seq results are consistent with these previous reports. Auxin is low in the stem cell region but accumulates in organ primordia to trigger differentiation in the SAM of angiosperms, supporting the view that the role of auxin in promoting cell differentiation is shared between bryophytes and angiosperms.^81^ We also noticed expression of several key genes involved in meristem function in angiosperms during merophyte development, including orthologs of *CLV1*, *ERL*, *SHR*, *HD-ZipIII*, and other homeobox transcription factors.^5,40,43,47,52,53^ These genes likely represent ancient developmental toolkits that regulate the growth of differentiating cells to form complex, three-dimensional organs and tissues. Activation of these genes reflects the complex, three-dimensional development in gametophores.

Our analysis also revealed diverse molecular mechanisms regulating SAM between bryophytes and angiosperms. In angiosperms, *KNOXI* genes promote cytokinin biosynthesis, which activates *WUSCHEL* (*WUS*), a master regulator of pluripotent stem cell proliferation in the SAM.^9^ However, in *P. patens*, expression of *KNOXI* orthologs and *WOX* family genes is not specific to the SAM (Figure S6), consistent with previous reports on tissue specificity.^17,18^ These diversifications likely reflect distinct evolutionary trajectories between the SAM of bryophytes and angiosperms. Recent studies indicate that *KNOXI* functions exclusively in sporophyte development in *P. patens*, while *WOX* genes were likely recruited for SAM regulation after the diversification of seed plants (Figure 5J).^10,73,82^

Pluripotent stem cells are maintained at the tip of the SAM and continuously differentiate organs in both bryophytes and angiosperms.^10,14^ However, the bryophytes SAM contains a single pluripotent stem cell and forms in the gametophyte, whereas the angiosperms SAM possesses multiple stem cells and forms in the sporophyte.^8,10^ Despite these differences, we have demonstrated the conserved cytokinin-ESR pathway promoting stem cell identity, as well as striking similarities in cytokinin and auxin distribution between the SAM of bryophytes and angiosperms. These findings suggest that high cytokinin, low auxin, and downstream factors including ESR regulated by these phytohormones are ancient and fundamental components underlying pluripotency in SAM. During the early evolution of land plants, the gene regulatory network related to these components might have been co-opted from the gametophytic SAM to the sporophytic SAM. The intriguing point is that the localizations of these components are condensed at the single pluripotent stem cell in bryophytes. This contrasts the angiosperm’s SAM, which has multiple domains interacting with each other.^9^ Specifically, the interaction between the organizing center and central region of angiosperm’s SAM was likely acquired during the evolution of complex SAM to stably maintain the multiple stem cells. To elucidate how the *KNOXI*, *WOX*, and other factors have been recruited into the SAM regulation after the divergence of vascular plants may lead to a further understanding of principle and evolution of pluripotency in land plants.

## STAR Methods

### KEY RESOURCES TABLE

| REAGENT or RESOURCE | SOURCE | IDENTIFIER |
| --- | --- | --- |
| **Bacterial and virus strains** |  |  |
| Escherichia coli DH5 $\alpha$ strain | Widely distributed | N/A |
| Escherichia coli ccdB Survival 2 T1R strain | Thermo Fisher Scientific | Cat#A10460 |
| **Chemicals, peptides, and recombinant proteins** |  |  |
| BCD medium | Nishiyama et al. 2000<sup>83</sup> | N/A |
| BCDAT medium | Nishiyama et al. 2000<sup>83</sup> | N/A |
| Plant agar | Nacalai tesque | Cat#01028-85 |
| Driselase | ASKA Animal Health Co., Ltd. | N/A |
| Polyethylene glycol (PEG) 6000 | FUJIFILM Wako Chemicals | Cat#169-09125 |
| G418 disulphate | FUJIFILM Wako Chemicals | Cat#078-05961 |
| Blasticidin S | funakoshi | Cat#KK-400 |
| 6- benzylaminopurine (BAP) | Sigma-Aldrich | Cat#B3408-5G |
| Dimethyl Sulfoxide (DMSO) | FUJIFILM Wako Chemicals | Cat#049-07213 |
| X-Gluc | Bio Medical Science Inc. | Cat#BC-GLUC100 |
| Propidium iodide (PI) solution | Sigma-Aldrich | Cat#P4864 |
| percoll | Cytiva | Cat#17089102 |
| RNA later | Thermo Fisher Scientific | Cat#AM7020 |
| Recombinant RNase Inhibitor | Takara Bio Inc. | Cat#2313A |
| Complete protease inhibitor cocktail | Roche | Cat#11873580001 |
| Sucrose | FUJIFILM Wako Chemicals | Cat#196-00015 |
| Sodium chloride | FUJIFILM Wako Chemicals | Cat#191-01665 |
| HEPES | FUJIFILM Wako Chemicals | Cat#346-01373 |
| Spermine tetrahydrochloride | Tokyo Chemical Industry Co., Ltd. | Cat#P2950 |
| Spermidine phosphate | Tokyo Chemical Industry Co., Ltd. | Cat#S0068 |
| Triton X-100 | Sigma-Aldrich | Cat#X100-500ML |
| β-Mercaptoethanol | FUJIFILM Wako Chemicals | Cat#131-14572 |
| Magnesium chloride | FUJIFILM Wako Chemicals | Cat#136-03995 |
| Ethylenediamine-N,N,N',N'-tetraacetic<br>Acid Disodium Salt Dihydrate (2NA)<br>(EDTA) | FUJIFILM Wako Chemicals | Cat#345-01865 |
| 1% Purified BSA solution | New England Biolabs | Cat#B9001S |
| Dithiothreitol | FUJIFILM Wako Chemicals | Cat#048-29224 |
| 4',6-diamidino-2-phenylindole (DAPI) | Thermo Fisher Scientific | Cat#62247 |
| **Critical commercial assays** |  |  |
| Chromium Next GEM Single Cell 3'<br>GEM, Library & Gel Bead Kit v3.1 | 10x Genomics | Cat#PN-1000128 |
| Chromium Next GEM Chip G Single<br>Cell Kit | 10x Genomics | Cat#PN-1000127 |
| Single Index Kit T Set A | 10x Genomics | Cat#PN-1000213 |
| NEBNext poly(A) mRNA Magnetic<br>Isolation Module | New England Biolabs | Cat#E7490S |
| NEBNext Ultra II RNA Library Prep Kit<br>for Illumina | New England Biolabs | Cat#E7770S |
| NEBNext Single Cell/Low Input RNA<br>Library Prep Kit for Illumina | New England Biolabs | Cat#E6420S |
| AMPure XP Reagent | Beckman Coulter | Cat#A63881 |
| RNeasy plant mini kit | QIAGEN | Cat#74904 |
| FastGene Plasmid Mini kit | NIPPON genetics | Cat#FG-90402 |
| GO Taq colorless | Promega | Cat#M7132 |
| Prime STAR GXL DNA polymerase | Takara Bio Inc. | Cat#R050A |
| Gateway BP Clonase II Enzyme mix | Thermo Fisher Scientific | Cat311789020 |
| **Deposited data** |  |  |
| snRNA-seq | This paper | PRJNA1138952 |
| bulk RNA-seq (young gametophores<br>and protonema) | This paper | PRJNA1140844 |
| bulk RNA-seq (cytokinin treatment) | This paper | PRJNA1140825 |
| **Experimental models:**<br>**Organisms/strains** |  |  |
| Physcomitrium patens Gransden | Widely available | N/A |
| P. patens: proPp3c21_6170:NGG | This paper | N/A |
| P. patens: proPp3c12_13360:NGG | This paper | N/A |
| P. patens: proPp3c7_3350:NGG | This paper | N/A |
| P. patens: proPp3c3_34230:NGG | This paper | N/A |
| P. patens: proPp3c27_870:NGG | This paper | N/A |
| P. patens: proPp3c24_12780:NGG | This paper | N/A |
| P. patens: proPp3c20_21980:NGG | This paper | N/A |
| P. patens: proPpLOG2:NGG | This paper | N/A |
| P. patens: Ppesr1 #4 | This paper | N/A |
| P. patens: Ppesr1 #10 | This paper | N/A |
| P. patens: quadruple #63 | This paper | N/A |
| P. patens: quadruple #127 | This paper | N/A |
| P. patens: quintupule #5 | This paper | N/A |
| P. patens: quintuple #15 | This paper | N/A |
| **Oligonucleotides** |  |  |
| See Table S2 for Primers | This paper | N/A |
| **Recombinant DNA** |  |  |
| pPIG1b:NGGII | Ishikawa et al. 2011<sup>84</sup> | AB537478 |
| pPIG1b:NGGII_proPp3c21_6170 | This paper | N/A |
| pPIG1b:NGGII_proPp3c12_13360 | This paper | N/A |
| pPIG1b:NGGII_proPp3c7_3350 | This paper | N/A |
| pPIG1b:NGGII_proPp3c3_34230 | This paper | N/A |
| pPIG1b:NGGII_proPp3c27_870 | This paper | N/A |
| pPIG1b:NGGII_proPp3c24_12780 | This paper | N/A |
| pPIG1b:NGGII_proPp3c20_21980 | This paper | N/A |
| pPIG1b:NGGII_proPpLOG2 | This paper | N/A |
| pDONR221 | Thermo Fisher Scientific | Cat#12536-017 |
| pPpU6-sgRNA | Lopez-Obando et al. 2016<sup>85</sup> | N/A |
| pPpU6-sgRNA_PpESR1 | This paper | N/A |
| pPpU6-sgRNA_PpESR2 | This paper | N/A |
| pPpU6-sgRNA_PpESR3 | This paper | N/A |
| pPpU6-sgRNA_PpESR4 | This paper | N/A |
| pPpU6-sgRNA_PpESR5 | This paper | N/A |
| pPpU6-sgRNA_PpESR6 | This paper | N/A |
| pPpU6-sgRNA_PpESR7 | This paper | N/A |
| pPpU6-sgRNA_PpESR8 | This paper | N/A |
| pPpU6-sgRNA_PpESR9 | This paper | N/A |
| pAct-Cas9 | Lopez-Obando et al. 2016<sup>85</sup> | N/A |
| pBNRF | Menand et al. 2007<sup>86</sup> | N/A |
| **Software and algorithms** |  |  |
| Cutadapt (v4.1) | Martin 2011<sup>87</sup> | RRID:SCR_011841 |
| Trimmomatic (v0.39) | Bolger et al. 2014<sup>88</sup> | RRID:SCR_011848 |
| fastq-pair (v0.4) | J. A Edwards and R. A. Edwards 2019<sup>89</sup> | https://github.com/linsalrob/fastq-pair |
| HISAT2 (v2.2.1) | Kim et al. 2019<sup>90</sup> | RRID:SCR_015530 |
| featureCounts (v2.0.1) | Liao et al. 2014<sup>91</sup> | RRID:SCR_012919 |
| Galaxy | Afgan et al. 2018<sup>92</sup> | RRID:SCR_006281 |
| Cell Ranger (v6.0.1) | 10x Genomics | RRID:SCR_017344 |
| R (v4.2) | R Development Core Team | RRID:SCR_001905 |
| Seurat (v4.2.0) | Hao et al. 2021<sup>93</sup> | RRID:SCR_007322 |
| Monocle3 (v1.3.1) | Qiu et al. 2017<sup>94</sup> | RRID:SCR_018685 |
| SeuratWrappers (v0.3.0) | Cao et al. 2019<sup>95</sup> | RRID:SCR_022555 |
| leiden (v0.4.3) | Traag et al. 2019<sup>96</sup> | https://github.com/TomKellyGenetics/leiden |
| ComplexHeatmap (v2.13.1) | Gu et al. 2016<sup>97</sup> | RRID:SCR_017270 |
| TCC (v1.34.0) | Sun et al. 2013<sup>98</sup> | RRID:SCR_001779 |
| multcomp (v1.4) | Hothorn et al. 2008<sup>99</sup> | RRID:SCR_018255 |
| g:Profiler | Kolberg et al. 2023<sup>100</sup> | RRID:SCR_006809 |
| Phytozome (v13) | Goodstein et al. 2012<sup>101</sup> | RRID:SCR_006507 |
| Black Zen software | Carl Zeiss | RRID:SCR_018163 |
| ImageJ (v1.53a) | Schneider et al. 2012<sup>102</sup> | RRID:SCR_003070 |
| CRISPOR | Concordet and Haeussler 2018<sup>103</sup> | RRID:SCR_015935 |
| Geneious (v2021.2.2) | Dotmatics | RRID:SCR_010519 |
| FigTree (v1.4.4) | Rambaut 2018 | RRID:SCR_008515 |
| **Other** |  |  |
| Chromium Controller | 10x Genomics | N/A |
| Hemocytometer C-Chip | NanoEntek | Cat#DHC-F01 |
| Falcon 40 µm Cell Strainer | Corning | Cat#352340 |
| Falcon 70 µm Cell Strainer | Corning | Cat#352350 |
| CellTrics filters 10 µm | Sysmex | Cat#04-004-2324 |
| Zeiss LSM880 | Carl Zeiss | N/A |
| Leica M165FC | Leica Microsystems | N/A |
| Olympus BX51 | EVIDENT | N/A |

### RESOURCE AVAILABILITY

#### Lead contact

Further information and requests for resources and reagents should be directed to and will be fulfilled by the lead contact, Junko Kyozuka.

#### Materials availability

Plasmids and transgenic plants generated in this study will be available on request to the lead contact, Junko Kyozuka.

#### Data and code availability

All raw RNA-seq data generated in this study were deposited at NCBI’s Sequence Read Archive (SRA). The accession number for the raw snRNA-seq data is PRJNA1138952. The accession number for the raw bulk RNA-seq data from the protonema and young gametophore samples is PRJNA1140844. The accession numbers for the raw bulk RNA-seq data from the cytokinin-treated samples are PRJNA1140825. This study used codes described in Quantification and statistical analysis.

## EXPERIMENTAL MODEL AND SUBJECT DETAILS

All experiments were performed using the Gransden Wood strain (1962) of *Physcomitrium patens* (*P. patens*).^104^ Plants were cultured and propagated in 9 cm dishes overlayed with BCDAT medium containing 0.8 % agar (Nacalai Tesque) under continuous light at 25℃, and then transferred to media for analyses.

We used the pPIG1b:NGG vector to construct the promoter-reporter lines for the gametophore markers, axillary hair markers, meristem marker, and proPpLOG2:NGG.^84^ Approximately 3.5 kb of promoter region was amplified by PCR, and cloned into a SmaI site of pPIG1b:NGG vector using the SLiCE method.^105^

We conducted CRISPR/Cas9 mediated knock-out (KO) of *PpESR* genes following the method of Lopez-Obando et al. 2016 with some modifications.^85^ Target sites for gRNA with high specificity were screened using the web-based tool CRISPOR, and 20 bp target sites located in the AP2 domain-coding region were selected. Chemically synthesized DNA fragments containing the U6 promoter of *P. patens* and gRNA scaffolds were cloned into pDONR201 (Thermo Fisher Scientific) vector to generate the pPpU6-sgRNA vector. Then, the gRNA target sequence specific to the target genes was introduced through site-directed mutagenesis mediated by inverse PCR.

Polyethylene glycol (PEG) mediated transformation was performed for the transformation of *P. patens*, as described by Nishiyama et al. 2000.^83^ All vectors introduced into *P. patens* were propagated in *Escherichia coli* (*E. coli*) and purified using the FastGene Plasmid Mini Kit (Nippon Genetics Co., Ltd.). For the construction of promoter-reporter lines, purified vectors were linearized using the PmeI restriction enzyme (NEB), and 15 µg of products per construct were used for the PEG-mediated transformation. Stable transformants were obtained through two rounds of antibiotics based selection (100 mg/L Blastcidine S, funakoshi). Integration of the constructs to the genome was confirmed by PCR-based genotyping. More than two independent lines for each construct were obtained. For the generation of KO mutants of the *PpESR* genes, the vectors for Cas9 expression (pAct-Cas9), antibiotics-resistant gene expression (pBNRF), and gRNA expression (pPpU6-sgRNA) were mixed as follows: 5 µg of pAct-Cas9, 1 µg of pBNRF, and 5 µg of pPpU6-sgRNA (per targets). Plants transiently transformed with these constructs were obtained following a single round of antibiotic selection (100 mg/L G418, FUJIFILM Wako Chemicals). Genotyping was performed by Sanger sequencing following PCR amplification of genomic DNA fragments around the gRNA target site.

Primers used for vector construction and genotyping in this study are shown in Table S2.

## METHOD DETAILS

### Nuclei isolation for snRNA-seq, 10x Genomics library construction, and sequencing

Nuclear isolation was performed using a combination of techniques, including Somssich’s method for nuclear run-on assay, Sikorskaite’s method for nuclear isolation from apple, potato, and tobacco leaf tissues, and Lang’s method for chloroplast isolation from *P. patens*.^106 – 108^ In brief, fragmented protonemata were grown for 12 days on BCD medium plates between two cellophane membranes, and approximately 1.4 g of plant tissues containing young gametophores were collected. The fresh plant tissues were fixed in 20 mL of 1% (v/v) formaldehyde cooled on ice for 15 minutes. The plant material was then washed with ice-cold purified water and rapidly frozen in liquid nitrogen. The frozen plant material was then ground in a mortar cooled in liquid nitrogen, and the frozen powder was dissolved in 10mL of RNAlater (Thermo Fisher Scientific). This suspension was filtered sequentially through 70 μm and 40 μm cell strainers, and 30 mL of buffer A HEPES [250 mM Sucrose, 10 mM NaCl, 50 mM HEPES, 0.15 mM Spermine tetrahydrochloride, 0.5 mM Spermidine phosphate, 20 mM EDTA, 0.035% (v/v) β-Mercaptoethanol, 1 mM MgCl_2_, 1x cOmplete Protease Inhibitor Cocktail EDTA free] was added to the filtrate. The mixture was centrifuged at 1600 g for 10 minutes at 4°C using a swing rotor. The supernatant was carefully removed, and the pellet was resuspended in 10mL of buffer A HEPES containing 0.2% (v/v) Triton. Then, a density gradient was created by layering 2 mL each of buffer A HEPES Triton containing 40%, 30%, and 10% Percoll (Cytiva) in a 15 mL tube. 5 mL of the suspension was layered on top of the density gradient in two separate tubes, and the tubes were centrifuged at 1600 g for 30 minutes at 4°C using a swing rotor. The nuclei, which were concentrated in the 40% fraction, were collected and gently mixed into 45 mL of ice-cold PBS. The mixture was then centrifuged at 1600 g for 10 minutes at 4°C using a swing rotor. The supernatant was carefully removed, and the pellet was resuspended in 200 μL of PBS with 0.1% BSA (NEB) and RNase inhibitor (Takara). The concentration of nuclei in this suspension was determined by counting 10 μL of the suspension using a hemocytometer. According to the protocol of 10x Chromium (10x Genomics), the concentration of the suspension was adjusted to target approximately 5,000 cells and then loaded into an emulsion generation device (Chromium controller) for single-cell nuclear cDNA synthesis and barcoding. Library preparation was performed using the 10x Chromium Next GEM Single Cell 3ʹ Reagent Kits v3.1 (10x Genomics). The procedure followed the protocol provided with the kit. The prepared libraries were sequenced using DNB-seq (BGI), with each sample sequenced on one lane, generating paired-end 100 bp reads.

### Sample preparation for bulk RNA-seq, library construction, and sequencing

To verify that the snRNA-seq data, RNA-seq data was also obtained from bulk samples of the protonemata and the young gametophores for comparison. For the protonema bulk sample, fragmented protonemata were cultured on BCD medium plates overlayed with a cellophane membrane for 1 week. Approximately 500 mg of flesh protonema tissues were collected per sample in three replicates and immediately frozen in liquid nitrogen. RNA was extracted using the RNeasy Plant Mini Kit (Qiagen). The obtained total RNA (100 ng) was used to prepare libraries using NEBNext Ultra II RNA Library Prep Kit for Illumina (NEB). For the bulk sample of the young gametophores, fragmented protonemata were cultured between two cellophane membranes on BCD medium plates for 12 days until the plants formed gametophores vigorously. Gametophores were harvested using microscissors to collect 200 gametophores for each sample in 4 replicates, and RNA was extracted using the RNeasy Plant Mini Kit. The extracted total RNA (0.8 ng) was used to prepare RNA-seq libraries using NEBNext Single Cell/Low Input RNA Library Prep Kit for Illumina (NEB). Sequencing (150 bp, paired ends) was performed using the DNB-seq (BGI).

For the bulk RNA-seq on the cytokinin-treated plants, a small amount of protonema tissue (approximately 0.5 mm in diameter) was inoculated between two cellophane membranes and cultured for 2 weeks on solid BCD medium plates. Around 15 colonies were collected per sample in four replicates, and transferred to liquid BCD medium containing 100 nM 6-benzylaminopurine (BAP) or solvent (DMSO). The liquid medium culture was conducted for 3 hours, and then the fresh plant tissues were immediately frozen in liquid nitrogen. RNA was extracted using the RNeasy Plant mini kit. RNA library construction and sequencing (100 bp, paired ends) were performed using the DNB-seq pipeline (BGI).

### Observation of plant growth

For the phenotypic analyses of gametophore formation and colony growth in the *Ppesr* mutants, a small amount of protonema tissue (approximately 0.5 mm in diameter) was inoculated between two cellophane membranes and cultured on solid BCD medium plates. Cytokinin response of *Ppesr* mutants was examined using solid BCD medium containing 100 nM BAP or solvent (DMSO).

### Histochemical GUS activity assay

GUS staining was conducted following the method of Aoyama et al. (2012) with minor modifications.^74^ Plant tissues were fixed with a fixation solution [0.2% (w/v) MES (pH 5.6), 0.3% (v/v) formalin, 0.3 M mannitol] at room temperature for 10 minutes. After washing with 50 mM NaH_2_PO_4_ (pH 7.0), the fixed tissues were vacuum-infiltrated with a substrate solution [50 mM NaH_2_PO_4_ (pH 7.0), 0.5 mM 5-bromo-4-chloro-3-indolyl β-D-glucuronide (X-Gluc), 0.5 mM K_3_Fe(CN)_6_, 0.5 mM K_4_Fe(CN)_6_, and 0.05% (v/v) Triton X-100] for 30 minutes and then stained at 37°C. After staining, the tissues were fixed with 5% (v/v) formalin for 10 minutes and subsequently immersed in 5% (v/v) acetic acid for 10 minutes. The stained and fixed tissues were dehydrated with an ethanol series. The tissues were then immersed in a chloral hydrate solution [66% (w/w) chloral hydrate, 8% (w/w) glycerol] at 4°C overnight for clearing before imaging.

### Microscopy

The GFP signal of the promoter GFP lines was observed using a confocal laser scanning microscope (Zeiss LSM880). Cell walls were stained with 10-50 mg/L of propidium iodide (PI) prior to observation. An LD LCI Plan-Apochromat 40x/1.2 Imm Korr DIC M27 or Plan-Apochromat 20x/0.80 M27 objective lens was used. A laser wavelengths of 543 nm was used for PI staining detection and 488 nm for GFP detection. For DAPI staining of plant tissues, fixation was done at room temperature for 30 minutes with fixing solution [10% (v/v) formaldehyde, 50 mM NaH_2_PO_4_], followed by washing with 50 mM NaH_2_PO_4_ (pH=7.0). The samples were then stained with DAPI solution [1 mg/L DAPI, 0.05% (v/v) TritonX-100] at room temperature for 20 minutes, washed again with 50 mM NaH_2_PO_4_ (pH=7.0), and imaged with LSM880. A Plan-Apochromat 20x/0.80 M27 objective lens was used, with a laser wavelength of 405 nm. The 3D reconstruction of z-stack images was done using the ZEN black software (Carl Zeiss). For imaging of the isolated nuclei, the nuclear suspension was supplemented with a 100 mg/L DAPI solution diluted 100 times. The observation was performed using an Olympus fluorescence microscope BX51 equipped with an Olympus DP71 (EVIDENT) camera. An UPlanFl 40× objective lens was used. GUS-stained samples were observed with the Olympus BX51 equipped with an Olympus DP71 camera, using a UPlanFl 40× objective lens. The plant growth phenotype of the *Ppesr* mutants was observed using a stereo microscope, M165FC (Leica Microsystems).

## QUANTIFICATION AND STATISTICAL ANALYSES

### Mapping, nucleus counting, and clustering of snRNA-seq data

All data from different replicates were processed separately until the data integration step in the clustering. After the sequencing of libraries, filtering based on quality scores was performed on the Read2 fastq files containing mRNA sequences from the obtained paired-end sequences. Sequences were processed using the Galaxy platform (https://usegalaxy.org/) as follows: First, TSO oligo sequences were removed using the Cutadapt, followed by removing low-quality base sequences using the SLIDINGWINDOW function of the Trimmomatic (window size = 4, QS>=30). Subsequently, sequences shorter than 15 bp were removed using the MINLEN function. Then, the filtered Read2 files were re-paired with the Read1 files containing cell barcodes using the fastq-pair, which were then used for mapping against the *P. patens* genome. For reference files used for mapping, a nuclear genome sequence file and an annotation file of *P. patens* (v3.3) were obtained from the Phytozome database (https://phytozome-next.jgi.doe.gov/). Chloroplast and mitochondrial genome sequence files and their annotation files were obtained from the EnsemblPlants database (https://plants.ensembl.org/index.html). The sequence files of nuclear genome, plastid, mitochondrion, and their annotation files were integrated into a single genome file and a single annotation file, respectively. Then, a reference file for mapping and nucleus counting was created using the “cellranger mkref” function of the Cell Ranger software (10x Genomics). Mapping and detection of nuclei were performed using the “cellranger count” function with the created reference file and the re-paired sequence data after filtering. The execution of the “cellranger count” was carried out on the supercomputer BIAS at the National Institute for Basic Biology using a singularity container of the Cell Ranger v6.0.1.

All downstream analyses after the mapping and counting were conducted in the R software. Dimensional reduction and clustering were performed using the Seurat package. In brief, Seurat objects were generated for each sample from the Cell Ranger outputs. Then, low-quality nuclei were removed by selecting only nuclei with unique molecular identifer (UMI) counts between 500 and 2500, gene counts between 400 and 2000, chloroplast gene UMIs less than 5%, and mitochondrial gene UMIs less than 1%. Subsequently, the Seurat object files from each sample were integrated into a single file, and the dimension reduction was performed using only genes on the nuclear genome. Batch correction with a “LogNormalize” method was applied in the integration step. An uniform manifold approximation and projection (UMAP) algorithm was used for dimension reduction, with 2000 variable genes and 50 principal components. A Leiden clustering was performed with the resolution value 0.005.

### Identification of gametophore marker genes from public RNA-seq data

We selected candidates of gametophore tissue-specific marker genes based on publicly available data from two previous studies that performed RNA-seq analyses using laser microdissection to isolate the gametophore apical cells and the protonema apical cells.^13,22^ Initially, genes significantly upregulated in the gametophore apical cells were selected.^13^ From these genes, those showing significant downregulation in the protonema apical cells were further selected.^22^ Seven genes with an expression level above 1000 RPM in the gametophore apical cells and with a Log2 fold change value (the gametophore apical cell samples vs whole plant body samples) of 3 or higher were used as marker genes. As a result of expression analyses using promoter-reporter lines, four gametophore marker genes, two axillary hair marker genes, and one meristem marker gene were identified.

### Validation of snRNA-seq by bulk RNA-seq

Bulk RNA-seq data from protonema and young gametophores were mapped to the *P. patens* genome separately (v3.3 genome, obtained from the Phytozome) using the HISAT2 package. The featureCounts package was used to generate read count data of all coding genes in the *P. patens* genome with the default parameter settings for paired-end reads. These processing were performed on the Galaxy platform. Checking variance between the samples and detection of DEGs were performed in the R software. Variance of the samples was examined by a principal component analysis (PCA) using the “prcomp” function following L1 normalization for the each sample. DEG detection was performed using the “estimateDE” function of the TCC package (FDR is 0.05 and a test method is “edger”) after TMM normalization. To check a correlation between the bulk RNA-seq data and the snRNA-seq data, firstly UMI counts of each gene in the snRNA-seq data were summed up separately in the protonema cell cluster (Cluster 1 and 3) and the gametophore cell cluster (Cluster 2 and 5). Spearman’s rank correlation coefficients between these pseudobulk UMI count data and the bulk RNA-seq count data were calculated.

### Identification of cluster-specific genes and pseudotime analysis

Specific genes expressed in each cluster were identified using the “FindAllMarkers” function of the Seurat with parameter settings as follows: logfc.threshold = 1, min.pct = 0.1, and p_val_adj <= 0.00001. The pseudotime analysis was performed using the Monocle3 package. First, the Seurat object after clustering was converted into a cds object readable by Monocle3 using the SeuratWrappers package. The imported UMAP plot into the Monocle3 was preprocessed by the Leiden clustering (resolution = 0.0005). Prediction of cell fate trajectory was conducted using the "learn_graph" function of Monocle3. Afterwards, target nuclei were narrowed down to those around the trajectory from protonema to gametophore cell clusters using the “choose_cells” function. A root node was set at a terminus of the protonema cluster side to calculate the pseudotime. Subsequently, genes showing significant expression changes along the pseudotime were detected using the “graph_test” function (“principal_graph” was used for neighbor graph, and q value threshold was set to 0.05). Patterns of gene expression changes detected were normalized by calculating the z-score and further smoothed using the “smooth.spline” function of the R software. Then, the genes were classified into 10 clusters by k-means clustering using the ComplexHeatmap package.

### GO enrichment analysis

GO enrichment analyses were performed using the g:GOSt function of the g:Profiler database (https://biit.cs.ut.ee/gprofiler/gost) with default parameter settings. Only “driving terms” were used for interpretation of results to reduce redundancy of similar GO terms.

### Identification of cytokinin response

Mapping, read counts, checking sample similarities, and DEG detection of the bulk RNA-seq data from the cytokinin-treated samples and the control samples were conducted following the same procedures as with the bulk RNA-seq data from protonema and young gametophore samples described above.

### Phylogenetic analysis

Amino acid sequences of a set of homologs encoding the AP2/ERF domain were obtained based on BlastP searches using *Arabidopsis thaliana* ESR1/DRN as a query from the Phytozome database.

Target plant species were *Arabidopsis thaliana*, *Oryza sativa*, *Ceratopteris richardii*, *Selaginella moellendorffii*, *Marchantia polymorpha*, and *Physcomitrella patens*. The collected amino acid sequnces were aligned using MUSCLE and the conserved regions of the AP2/ERF domain were extracted for the following phylogenetic analysis. The phylogenetic tree was reconstructed based on Neighbor-Joining method with 10,000 times bootstrap replicates. The alignment, extraction, and phylogeny reconstruction were performed using the Geneious software. The phylogenetic tree was visualized using the FigTree software.

### Quantification of plant growth phenotype

The number of gametophores and protonema cell length were determined from pictures taken by a stereo microscope (M165FC, Leica), using the ImageJ software. Measurement of colony area was conducted as follows: A picture of a colony taken by the Leica M165FC was imported to ImageJ and the "minimum" filter (radius setting is 20 pixels) was applied following the conversion to the grayscale image. The filtered image was binarized using the "Threshold" function with default settings, and the area of the colony region was measured by the "Analyze particles" function. The number of cell divisions of the protonemal apical cell was counted from the overlayed semi-transparent images from day 6 and day 9 pictures of protonema taken by the Leica M165FC. The honestly significant difference (HSD) test was performed to evaluate the statistical significance using the multcomp package in the R software.

## Acknowledgments

We thank Dr. Fabien Nogué (INRAE, France) for providing pAct-Cas9 and pBNRF vectors, and detailed instructions for genome editing in *P. patens*. Computational resources to run the Cell Ranger software were provided by the Data Integration and Analysis Facility, National Institute for Basic Biology. We acknowledge funding supports from Japan Society for the Promotion of Science (JSPS) KAKENHI (23H05409, 20H05684, 17H06475, 17H06472, 20J20812 and 23K19362).

## Author contributions

Conceptualization: JK, YH ; Methodology: YH, NH, KB, AH, AM, MH ; Investigation: YH ; Visualization: YH ; Funding acquisition: JK, MH, YH ; Project administration: JK ; Supervision: JK ; Writing – original draft: JK, YH ; Writing – review & editing: JK, YH

