## Supplementary material for "snRNA-seq analysis of the moss *Physcomitrium patens* reveals a conserved cytokinin-ESR module promoting pluripotent stem cell identity": Figures S1 to S6

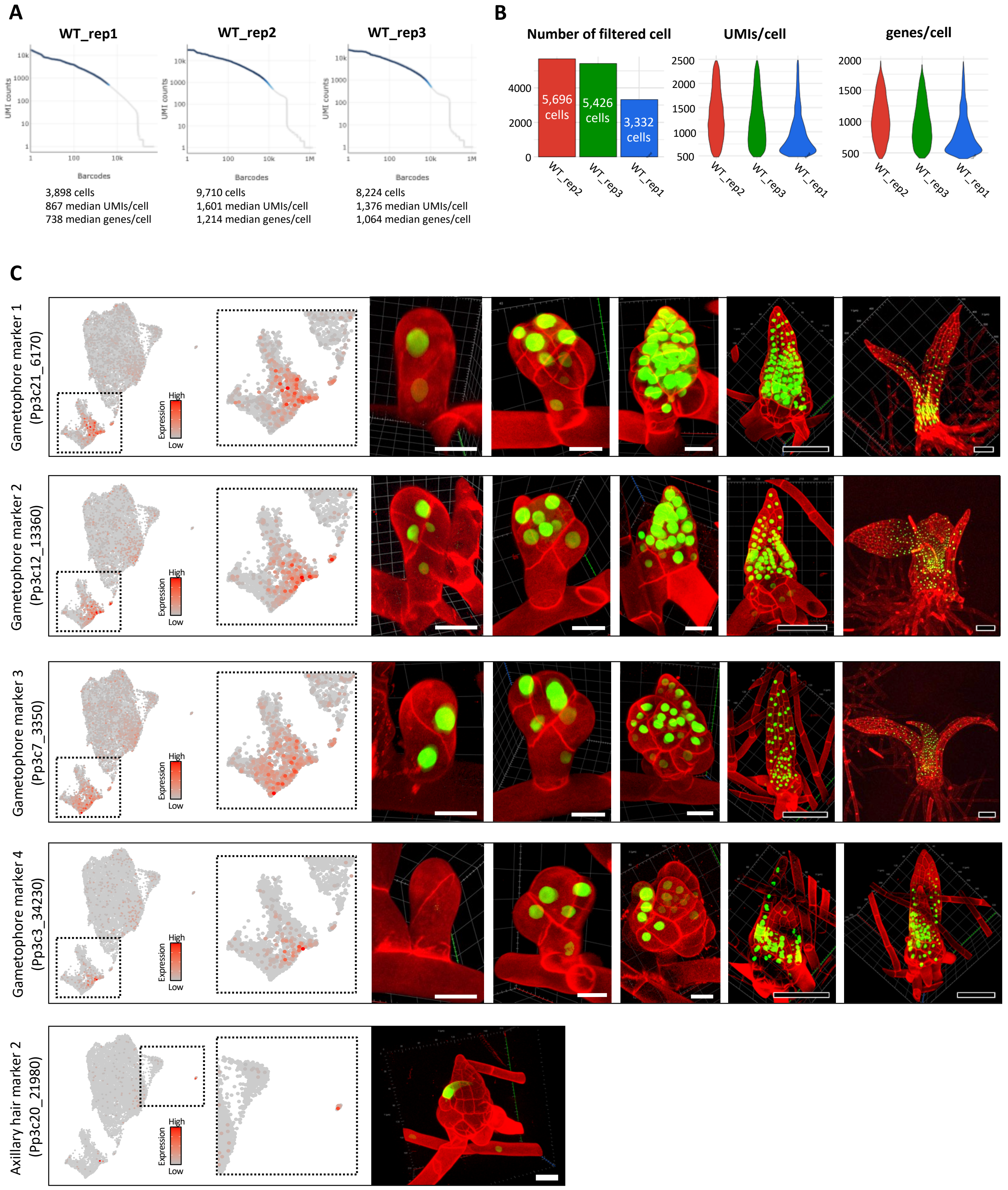


**Figure S1. Basic statistics of snRNA-seq data and expression patterns of marker genes**

(A) Barcode-rank plot and summary of the Cell Ranger outputs. Dark blue lines are barcodes identified as nuclei by the Cell Ranger program.

(B) Basic statistics of the dataset after the filtration of low-quality nuclei.

(C) Expression patterns of the gametophore markers and an axillary hair marker. Expression patterns in snRNA-seq are shown on the left, and the expression patterns of promoter GFP lines are shown on the right. Green and red signals indicate GFP and cell walls stained by propidium iodide, respectively. White scale bars, 20 µm. Black scale bars, 100 µm.

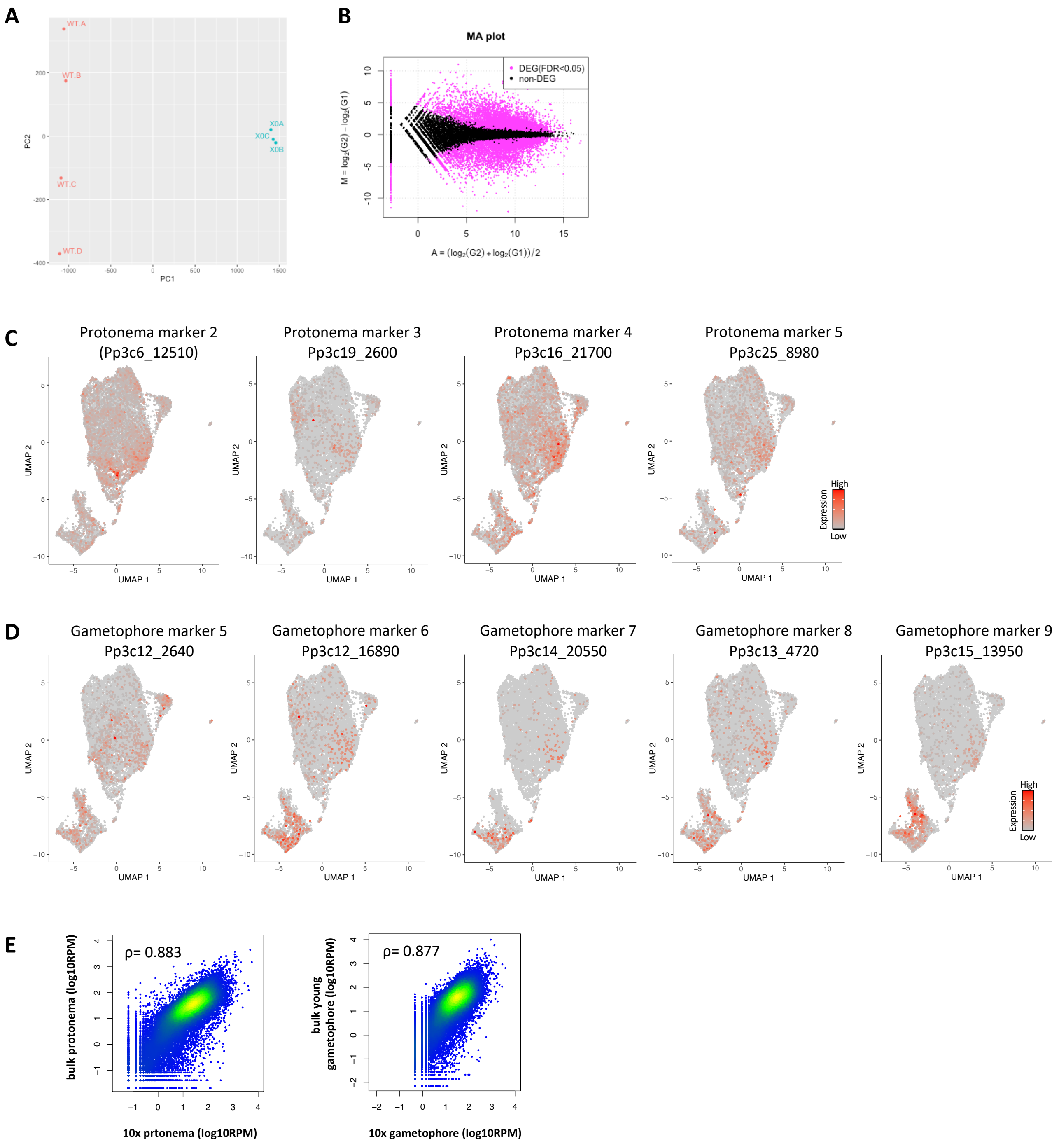


**Figure S2. Validation of snRNA-seq data by bulk RNA-seq**

(A) Principal component analysis (PCA) of bulk RNA-seq data from young gametophores and protonemata. WT.A to WT.D indicate the young gametophore samples, and X0A to X0C indicate the protonema samples.

(B) MA-plot of bulk RNA-seq data from young gametophores and protonemata. Each dot presents genes. Magenta-colored dots indicate the differentially expressed genes between the young gametophore and protonema samples (FDR = 0.05).

(C) Expression patterns of protonema marker genes identified from bulk RNA-seq. Protonema marker genes were selected from the top 5 highly expressed genes within the 100 genes that showed the lowest q value.

(D) Expression patterns of gametophore marker genes identified from bulk RNA-seq. The gametophore marker genes were selected from the top 5 highly expressed genes within the 100 genes that showed the lowest q value.

(E) Correlation between snRNA-seq data and bulk RNA-seq transcriptome data. Each dot indicates the expression level of a gene in the snRNA-seq and bulk RNA-seq. Correlation in the gametophore data and protonema data are shown. Spearman's rank correlation coefficients (ρ) were calculated.

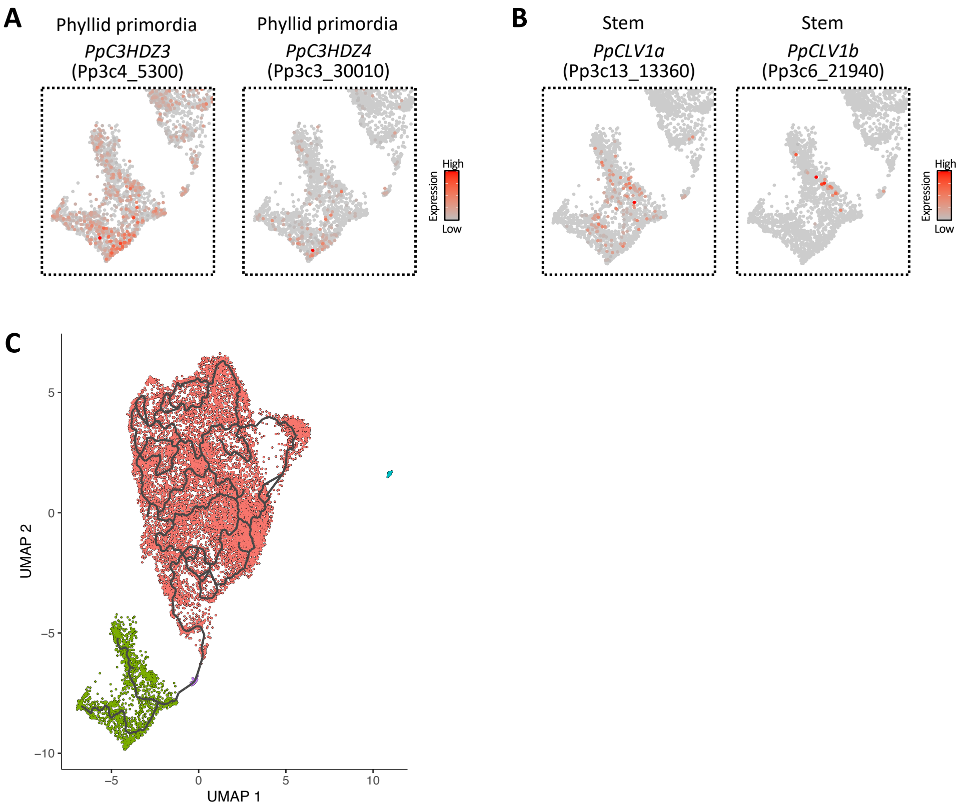


**Figure S3. Expression patterns of gametophore tissue marker genes and trajectory detection**
(A and B) Expression pattern of the known genes around the gametophore cluster. The genes known to be expressed in phyllid primordia (A) and stem tissue (B) are shown.
(C) Trajectory detected by the "learn_graph" function of the Monocle3 software.

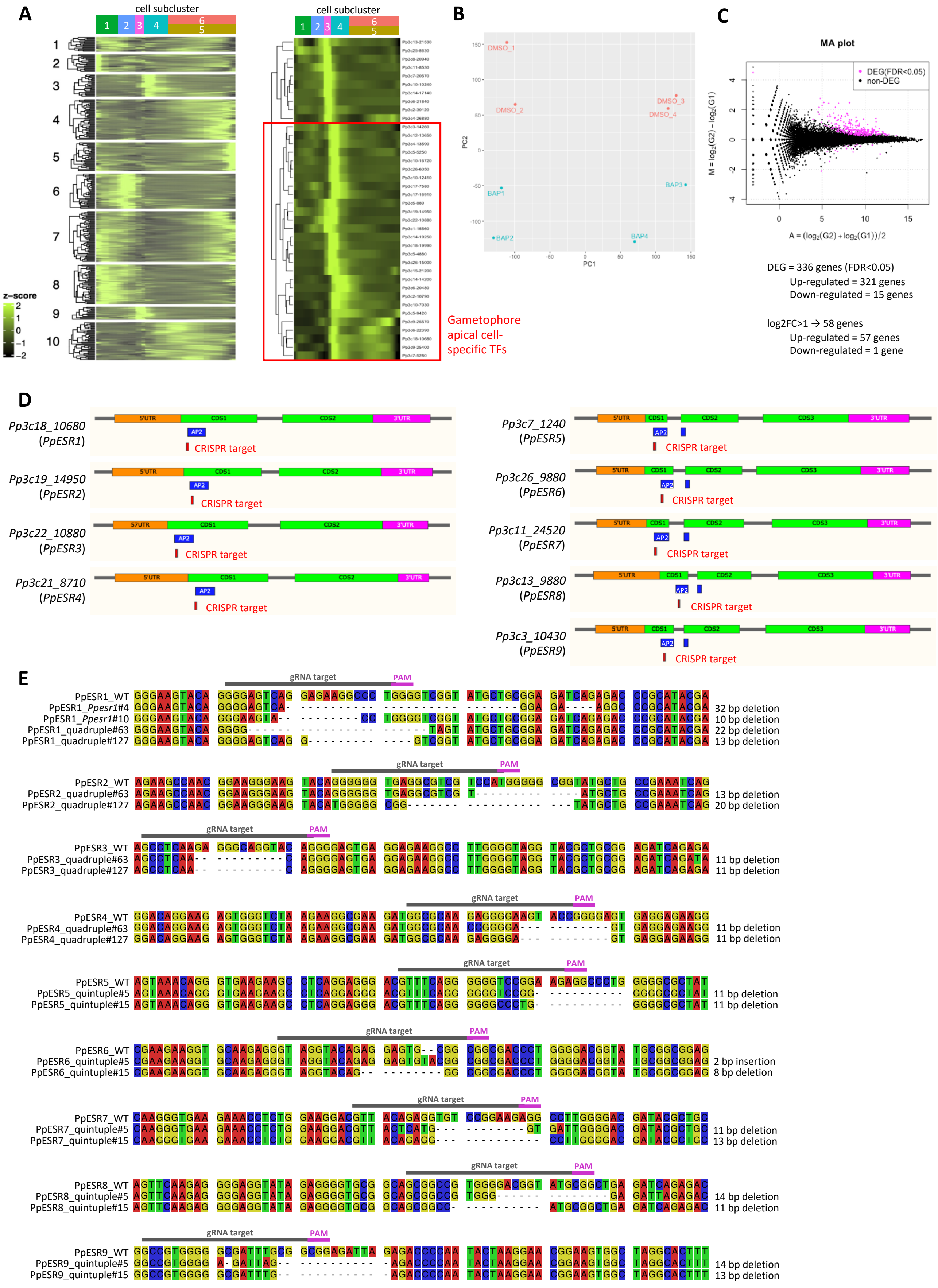
A
cell subcluster
1 2 3 4 6 5
1 2 3 4 5 6 7 8 9 10
z-score
2 1 0 −1 −2
Gametophore apical cell-specific TFs
B
DMSO_1 DMSO_2 DMSO_3 DMSO_4
BAP1 BAP2 BAP3 BAP4
PC1
PC2
C
MA plot
DEG(FDR<0.05)
non-DEG
M = log2(G2) − log2(G1)
A = (log2(G2) + log2(G1))/2
DEG = 336 genes (FDR<0.05)
Up-regulated = 321 genes
Down-regulated = 15 genes
log2FC>1 → 58 genes
Up-regulated = 57 genes
Down-regulated = 1 gene
D
Pp3c18_10680 (PpESR1)
Pp3c19_14950 (PpESR2)
Pp3c22_10880 (PpESR3)
Pp3c21_8710 (PpESR4)
Pp3c7_1240 (PpESR5)
Pp3c26_9880 (PpESR6)
Pp3c11_24520 (PpESR7)
Pp3c13_9880 (PpESR8)
Pp3c3_10430 (PpESR9)
5'UTR CDS1 CDS2 CDS3 3'UTR AP2
CRISPR target
E
gRNA target
PAM
PpESR1_WT
PpESR1_Ppesr1#4 32 bp deletion
PpESR1_Ppesr1#10 10 bp deletion
PpESR1_quadruple#63 22 bp deletion
PpESR1_quadruple#127 13 bp deletion
PpESR2_WT
PpESR2_quadruple#63 13 bp deletion
PpESR2_quadruple#127 20 bp deletion
PpESR3_WT
PpESR3_quadruple#63 11 bp deletion
PpESR3_quadruple#127 11 bp deletion
PpESR4_WT
PpESR4_quadruple#63 11 bp deletion
PpESR4_quadruple#127 11 bp deletion
PpESR5_WT
PpESR5_quintuple#5 11 bp deletion
PpESR5_quintuple#15 11 bp deletion
PpESR6_WT
PpESR6_quintuple#5 2 bp insertion
PpESR6_quintuple#15 8 bp deletion
PpESR7_WT
PpESR7_quintuple#5 11 bp deletion
PpESR7_quintuple#15 13 bp deletion
PpESR8_WT
PpESR8_quintuple#5 14 bp deletion
PpESR8_quintuple#15 11 bp deletion
PpESR9_WT
PpESR9_quintuple#5 14 bp deletion
PpESR9_quintuple#15 13 bp deletion

38

**Figure S4. Identification of gametophore apical cell-specific TFs and generation of *Ppesr* loss-of-function mutants**

(A) Expression patterns of transcription factors that showed significant expression changes along pseudotime from protonema to gametophore. k-means clustering was performed and genes included in group 3 and 9 were selected as the candidates for gametophore apical cell-specific TFs (the left heatmap). The candidate TFs were further classified by k-means clustering, and the lower biggest group, showing the highest expression level at the gametophore apical cell position and was identified as the gametophore apical cell-specific TFs (the right heatmap).

(B) Principal component analysis (PCA) of bulk RNA-seq data from the BAP treatment experiments. "DMSO_1" to "DMSO_4" are the control samples, and "BAP_1" to "BAP_4" are the BAP-treated samples.

(C) MA-plot of bulk RNA-seq data from the BAP-treated experiments. Each dot presents genes. Magenta colored dots indicate the differentially expressed genes between the control and BAP-treated samples (FDR = 0.05).

(D) Gene structures of *PpESR* genes. The CRISPR/Cas9 target sites were designed inside of the conserved AP2 domain.

(E) Mutation pattern of *Ppesr* mutants generated by the CRISPR/Cas9 system examined by the Sanger sequencing. All mutations cause frame-shift leading to premature arrest by a stop codon.

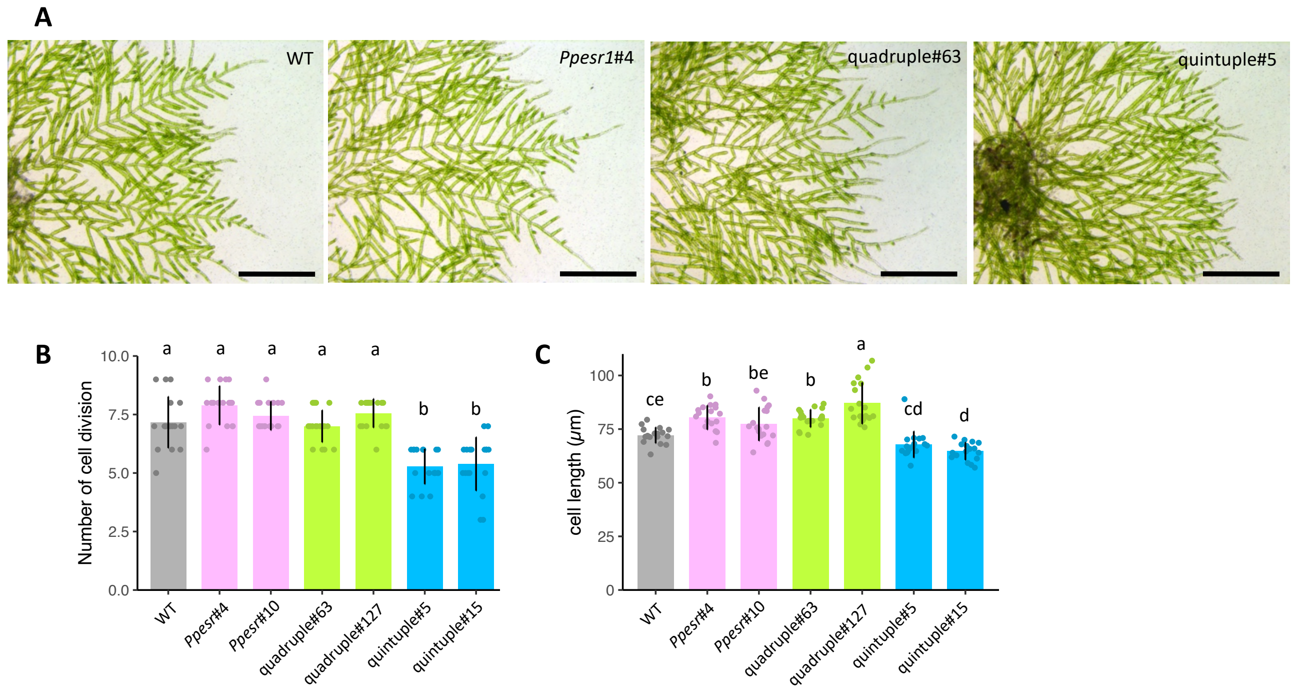


**Figure S5. Detailed phenotypes of *Ppesr* mutants in protonemata**

(A) Bright-field images of protonemata in WT and *Ppesr* mutants. Plates were cultured for 13 days

between two cellophane membranes on solid BCD medium. Scale bars, 500 µm.
(B and C) Phenotypes of *Ppesr* mutants in protonemal growth. Number of cell divisions in the caulonema apical stem cell from day 6 to day 9. Protonemata were grown between two cellophane membranes on solid BCD medium (B). Caulonemal cell length at day 9 when grown between two cellophane membranes on solid BCD medium (C). Statistical significance was assessed using the HSD test (n =30, $p < 0.05$).

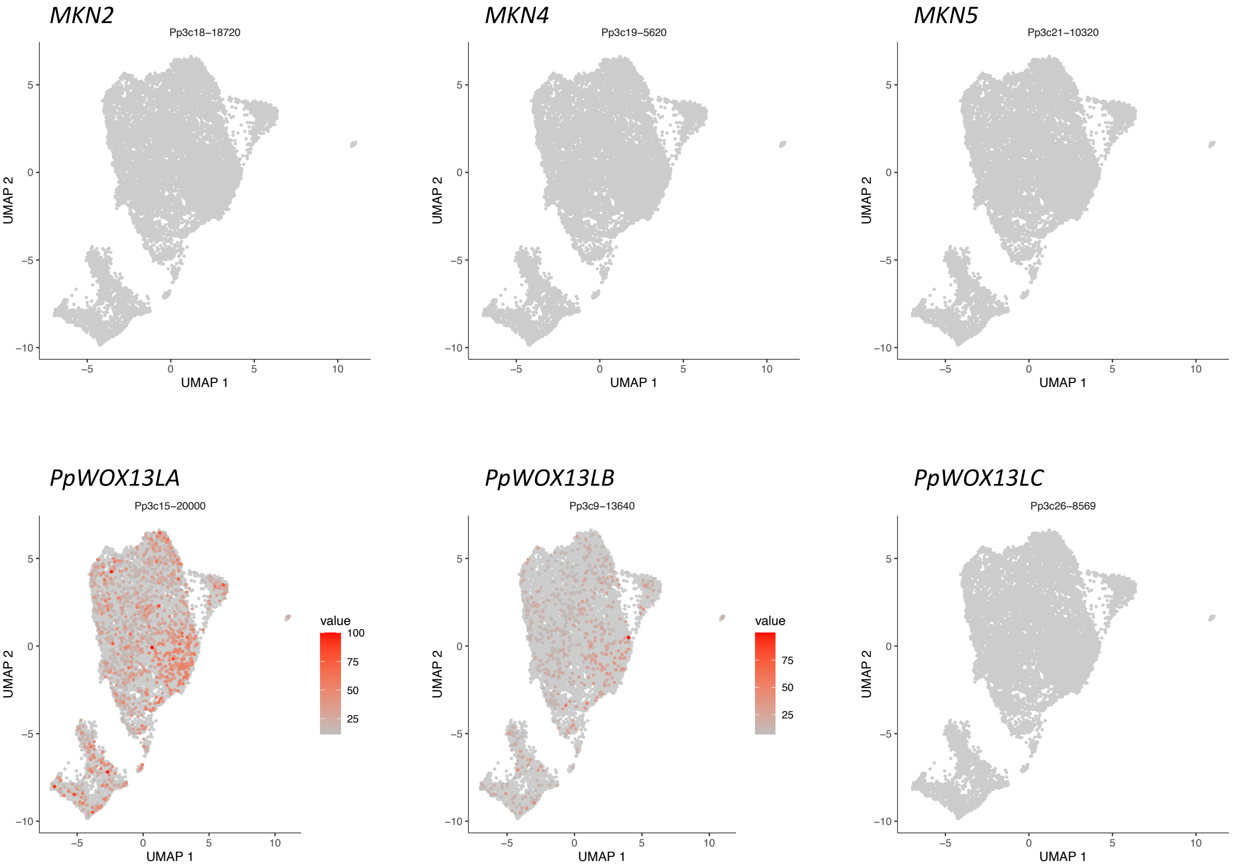


**Figure S6. Expression patterns of selected genes related to key factors in SAM of angiosperms**
